# Fasting Induces a Highly Resilient Deep Quiescent State in Muscle Stem Cells via Ketone Body Signaling

**DOI:** 10.1101/2022.01.04.474961

**Authors:** Daniel I. Benjamin, Pieter Both, Joel S. Benjamin, Christopher W. Nutter, Jenna H. Tan, Jengmin Kang, Leo A. Machado, Julian D. D. Klein, Antoine de Morree, Soochi Kim, Ling Liu, Hunter Dulay, Ludovica Feraboli, Sharon M Louie, Daniel K Nomura, Thomas A. Rando

## Abstract

Short-term fasting is beneficial for the regeneration of multiple tissue types. However, the effects of fasting on muscle regeneration are largely unknown. Here we report that fasting slows muscle repair both immediately after the conclusion of fasting as well as after multiple days of refeeding. We show that ketosis, either endogenously produced during fasting or a ketogenic diet, or exogenously administered, promotes a deep quiescent state in muscle stem cells (MuSCs). Although deep quiescent MuSCs are less poised to activate, slowing muscle regeneration, they have markedly improved survival when facing sources of cellular stress. Further, we show that ketone bodies, specifically β-hydroxybutyrate, directly promote MuSC deep quiescence via a non-metabolic mechanism. We show that β-hydroxybutyrate functions as an HDAC inhibitor within MuSCs leading to acetylation and activation of an HDAC1 target protein p53. Finally, we demonstrate that p53 activation contributes to the deep quiescence and enhanced resilience observed during fasting.

## INTRODUCTION

Dietary restriction (DR) robustly improves healthspan and lifespan in multiple model systems (Lee et al., 2006; McCay et al., 1935, 1989; Weindruch and Walford, 1982). Beneficial effects of dietary restriction have been observed at the organismal level, from simple eukaryotes to humans, and also at the single cell level in adult stem cell populations (Cheng et al., 2014; Jiang et al., 2000; Ravussin et al., 2015). These improvements are associated with cell and tissue functional maintenance and a delay or even prevention of age-related pathologies such as cognitive decline, inflammation, and cancer (Catterson et al., 2018; Colman et al., 2014; Weindruch et al., 1982). Commonly studied paradigms of dietary restriction are fasting and caloric restriction (CR). Fasting can lead cells and tissues to enter a protected state in which they become highly resistant to environmental stresses and toxicity (Mitchell et al., 2010; Raffaghello et al., 2008). For example, preoperative fasting lessens hepatic damage resulting from ischemia/reperfusion injury (Mitchell et al., 2010). In some contexts, fasting or CR has been shown to accelerate tissue regeneration. For example, intestinal epithelial and hepatic tissue regeneration after acute damage have been reported to improve after intense CR (Ramaiah et al., 1998; Yousefi et al., 2018).

Fasting affects a diverse population of stem and progenitor cells, frequently ameliorating various stem cell aging phenotypes (Mana et al., 2017). For example, fasting increases the ability of intestinal stem cells (ISCs) from young and aged mice to form intestinal organoids (Mihaylova et al., 2018). The extent of this functional improvement positively correlates with the duration of the fast and is dependent on ISC fatty acid oxidation. Similarly, the age-associated decrease in expression of the muscle stem cell (MuSC)-specific transcription factor Pax7 can be restored to youthful levels after multiple rounds of a short term calorically restrictive diet designed to mimic fasting (Brandhorst et al., 2015). This same fasting-mimicking diet (FMD) was also shown to rescue the decline in mesenchymal stem and progenitor cell number within bone marrow that is observed with aging (Brandhorst et al., 2015). Likewise, fasting or FMD promotes self-renewal of hematopoietic stem cells (HSCs), increases HSC number, and protects HSCs against cytotoxic agents (Brandhorst et al., 2015; Cheng et al., 2014). In some instances, the affected stem cells may even rejuvenate their associated tissue in response to DR. For example, periodic fasting corrects the myeloid bias observed in aged hematopoiesis (Cheng et al., 2014).

Adult stem cells are critically important for maintaining tissue integrity and for regenerating damaged tissue after injury. MuSCs are indispensable for muscle tissue repair (Lepper et al., 2011; Wang et al., 2014). Long term regenerative capacity of muscle is dependent on the ability of MuSCs to remain quiescent in the absence of injury, and inability to maintain quiescence results in defective muscle repair (Yue et al., 2017). In response to muscle injury, MuSCs enter the cell cycle, proliferate as myoblasts, and either self-renew to replenish the stem cell pool or fuse into nascent fibers to repair the injury (Lepper et al., 2011; Wang et al., 2014). In the absence of injury, quiescence is actively maintained and tightly regulated by a combination of signaling factors from the surrounding microenvironment and cell-intrinsic gene regulation, namely expression of cell cycle inhibitors and repression of cyclins, cyclin dependent kinases, and checkpoint kinases (Cheung and Rando, 2013; Fukada et al., 2007). For example, Notch signaling from the MuSC niche is known to be important in both maintenance of the quiescent state as well as self-renewal and reestablishment of quiescence following MuSC activation (Bjornson et al., 2012; Conboy et al., 2003; Mourikis et al., 2012; Wen et al., 2012). Quiescent MuSCs, like other quiescent stem cell populations, are characterized by stress resistance and low metabolic activity (Cheung and Rando, 2013). The effect of fasting on maintenance of this quiescent state, and on MuSC function in general, is unknown. The goal of this study was to comprehensively characterize the effect of fasting on muscle tissue repair and muscle stem cell function.

In this study, we show that fasting causes MuSCs to enter a state of deep quiescence that is characterized by delayed activation and enhanced resilience to nutrient, cytotoxic, and proliferative stress. In this state, MuSCs are functionally and transcriptionally less committed to a myogenic program and more stem-like as assessed by muscle regeneration and transplantation assays as well as RNA-seq analysis. Furthermore, increased MuSC resilience and delayed activation also results from feeding mice a ketogenic diet or injecting them with exogenous ketone bodies. Mechanistically, this deep quiescent state results from the inhibition of HDAC1 activity by increased levels of the primary circulating ketone body, β-hydroxybutyrate (BHB). We show, using pharmacological and genetic tools, that p53 activation downstream of HDAC1 inhibition is both necessary and sufficient to drive ketosis-induced deep quiescence. Altogether, this report highlights the novel finding that a metabolite produced endogenously during dietary restriction elicits a protective state in a stem cell population.

## RESULTS

### Fasting delays muscle regeneration, an effect that perdures after refeeding

To comprehensively characterize the effects of fasting on muscle regeneration, we injured the tibialis anterior (TA) muscle of the lower hindlimb in mice fasted for 0, 1, 2, or 2.5 days (**Figure 1A**). After 7 days of recovery, we isolated the muscles and examined regeneration histologically. We found that recovery, as measured by regenerating myofiber cross sectional area (CSA), progressively declined as a function of duration of fasting prior to injury (**Figures 1B** **and 1C)**. To explore the reversibility of this regenerative delay, we measured the extent of muscle repair in mice that had been fasted for 2.5 days and subsequently refed for 1, 2, 3, or 7 days prior to injury (**Figure 1D**). Intriguingly, we found that an impairment of regeneration persisted up to 3 days after refeeding, despite a complete return of body weight **(Figures 1E, 1F, and S1A)**. Importantly, one week of refeeding was able to restore muscle regeneration back to baseline. These findings indicate that fasting causes a transient state of impaired regenerative activity that persists days after refeeding.

**Figure 1.**
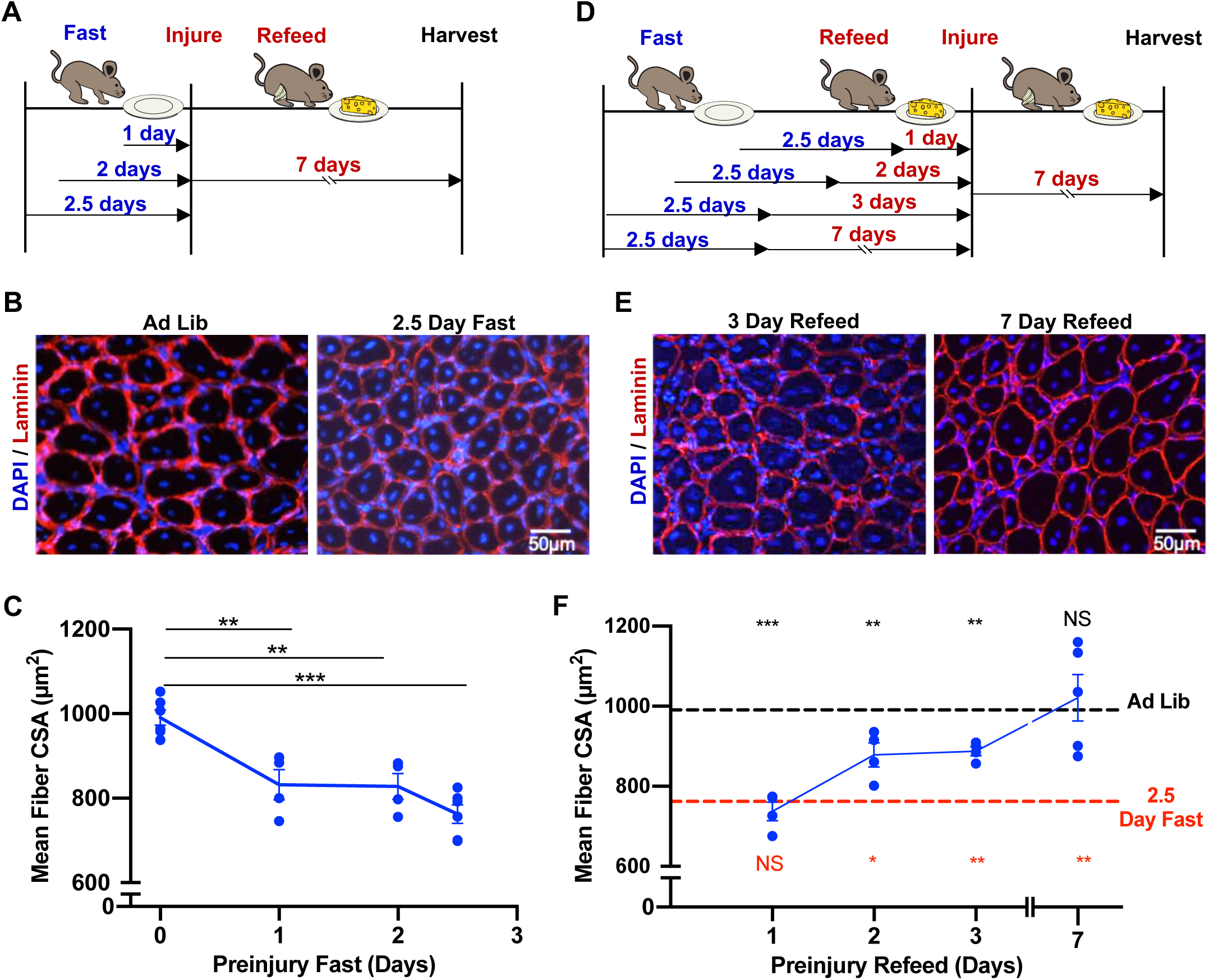
Fasting delays muscle regeneration, an effect that perdures after refeeding. **A**) Diagram of time course that depicts fasting durations before BaCl_2_ injury to the TA muscle, followed by refeeding and 7 days of recovery. **B**) Representative immunostaining of regenerating muscle from mice fed ad libitum or fasted for 2.5 days before injury. TA muscles were harvested 7 days after injury, sectioned, and stained for laminin. Nuclei were stained with DAPI. **C**) Quantification of regenerating muscle fiber cross-sectional area from mice fed ad libitum or fasted for 1, 2, or 2.5 days before injury as described in (A). **D**) Diagram of time course that depicts fasting and refeeding durations before BaCl_2_ injury to the TA muscle, followed by 7 days of recovery. **E**) Representative histology of regenerating muscle from mice that were fasted for 2.5 days and subsequently re-fed for 3 days or 7 days before injury. TA muscles were harvested 7 days after injury, sectioned, and stained for laminin. Nuclei were stained with DAPI. Quantification of regenerating muscle fiber cross-sectional area from mice fasted for 2.5 days then re-fed for 1, 2, 3, or 7 days before injury as described in (D). Error bars represent SEM. *p < 0.05; **p < 0.01; ***p < 0.001; ns, not significant. Black asterisks represent statistical comparisons to ad libitum. Red asterisks represent statistical comparisons to 2.5 days fasting. See also Figure S1.

### Fasting promotes a state of deep quiescence in muscle stem cells through ketosis

Because MuSCs are critically important for muscle regeneration after injury (Lepper et al., 2011; Wang et al., 2014), we next wanted to examine how short-term fasting might be affecting the function of the MuSCs themselves. After fasting a cohort of mice, we isolated quiescent MuSCs from the hindlimbs by FACS purification as previously described (Liu et al., 2015). We found that MuSCs from fasted mice were smaller than their control counterparts at the time of isolation (**Figure 2A**). Additionally, MuSCs from fasted mice exhibited a significant reduction in mitochondrial content, RNA content, and basal oxygen consumption compared with control MuSCs **(Figures 2B, S2A, and S2B)**. These cells were also significantly delayed in their time to enter S phase both after ex vivo activation as well as in response to in vivo injury (**Figures 2C** **and S2C)**. Therefore, MuSCs from fasted mice exhibit properties that are characteristic of cells in a state of deep quiescence (DQ) (Fujimaki et al., 2019; Kwon et al., 2017). In agreement with our muscle regeneration data, we found that this state of DQ persists up to 2 days after refeeding **(Figures S2D-S2F),** despite the return to baseline of body weight **(Figure S1A)**. These data suggest that DQ represents a stem cell state that can perdure for multiple days after the conclusion of fasting, and that this perdurance may explain the delay in muscle regeneration that we observed after fasting, even following 72 hours of refeeding.

**Figure 2.**
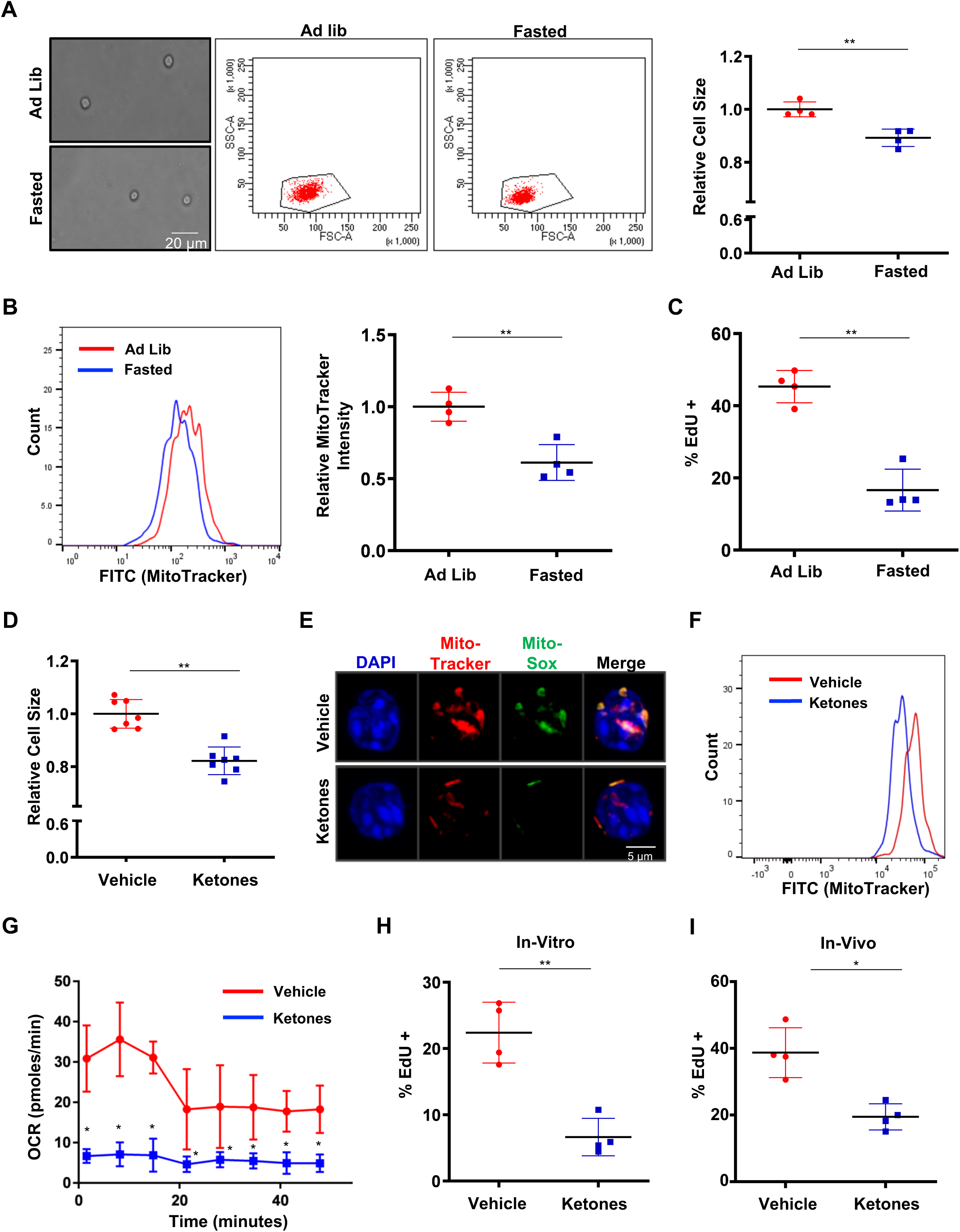
Fasting promotes a state of deep quiescence in muscle stem cells through ketosis. **A**) Representative brightfield images (left) and representative FACS plots (middle) of MuSCs harvested from fasted (60 hours) and ad lib fed mice. Right panel shows quantification of cell size based on forward scatter in FACS plots (n=4). **B**) Representative FACS plot (left) and quantification (right) of MitoTracker Green signal in MuSCs from fasted (60 hours) and ad lib fed mice (n=4). **C**) Quantification of EdU incorporation in MuSCs isolated from fasted (60 hours) and ad lib fed control mice after 48 hours in culture (n=4). **D**) Quantification of the cell size (based on forward scatter in FACS plots) of freshly isolated MuSCs from ketone body- or vehicle-treated mice. Mice were injected i.p. with ketone bodies (200 mg/kg BHB and 200 mg/kg acetoacetate) or vehicle three times per day for 1 week (n=4). **E**) Representative confocal microscopy images of freshly isolated MuSCs from ketone body- or vehicle-treated mice showing total mitochondrial staining with MitoTracker Deep Red as well as mitochondrially produced reactive oxygen species with MitoSox Green. Nuclei were stained with DAPI. **F**) Representative FACS plot of MitoTracker Green intensity in freshly isolated MuSCs from ketone body- or vehicle-treated mice (n=4). **G**) Seahorse assay results showing oxygen consumption rates (OCR) in freshly isolated MuSCs from ketone body- or vehicle-treated mice (n=3 per group). **H**) Quantification of EdU incorporation in MuSCs isolated from ketone body- or vehicle-treated mice and maintained in culture for 48 hours in the continuous presence of EdU (n=4). **I**) Cell cycle entry based on EdU incorporation of MuSCs from mice treated with ketone bodies or vehicle. Mice were injected with EdU (50 mg/kg) once daily for two weeks. Freshly isolated MuSCs were immediately fixed and stained for EdU. Error bars represent SEM. *p < 0.05; **p < 0.01; ***p < 0.001; ns, not significant. See also Figure S2.

Recent work has suggested that many of the beneficial effects of fasting (especially longer term fasting and caloric restriction) may be the result of the ketosis that accompanies the period of fasting (Veech et al., 2017). In order to assess whether or not ketosis, absent the stark energy imbalance seen with fasting, might be responsible for promoting this state of DQ, we fed mice an ad libitum ketogenic diet for 3 weeks to induce endogenous ketosis **(Figure S2G)**, and asked whether or not that might also induce a similar state of DQ in the MuSCs. We found that the MuSCs isolated from mice on the ketogenic diet exhibited the same hallmarks of the DQ state that we observed with fasting **(Figures S2H-S2K)**. Similarly, we detected a significant delay in muscle regeneration in mice that were fed a ketogenic diet **(Figure S2L and S2M).** Given that the ketogenic diet and fasting both involve drastic hormonal changes that accompany carbohydrate restriction, we next asked whether supplementation of an ad libitum chow diet with exogenous ketone bodies could also promote MuSC DQ. Mice were injected intraperitoneally with the two primary ketone bodies (BHB and acetoacetate) three times per day for 7 days in order to mimic the endogenous ketosis brought about by fasting and the ketogenic diet **(Figure S2N-Q)**. Unlike fasting or the ketogenic diet, exogenous ketone body supplementation resulted in no change in blood glucose levels or body weight **(Figures S2R and S2S)**. Similar to fasting and the ketogenic diet, exogenous ketone bodies also induced all of the hallmarks of DQ in MuSCs, including smaller size, less mitochondrial content, less oxygen consumption, less RNA content, and a slower rate of S phase entry **(Figures 2D-2H and S2T)**. We also found that the MuSCs from ketone-treated mice had a decreased propensity to break quiescence and enter S phase under homeostatic conditions in vivo (**Figure 2I**). Collectively, these data suggest that ketosis itself, and ketone bodies in particular, might be a primary driver of MuSC DQ.

### Ketosis promotes a cell cycle and myogenic signature consistent with deep quiescence

To probe the molecular changes underlying this ketone-induced deep quiescence (KIDQ), we sequenced the transcriptomes of freshly isolated MuSCs from mice that were injected with ketone bodies or vehicle control (**Figure 3A**). Consistent with our evidence that MuSCs from ketone body-treated mice exist in a deeper state of quiescence, gene ontology analysis revealed significant and robust changes in cell cycle genes, including downregulation of well-established proliferation genes such as the E2F family members, CDK2, and multiple members of the cyclin family of proteins (**Figures 3B** **and 3C**). We also observed an upregulation of genes involved in the inhibition of cell proliferation and growth, including CDKN1A (p21) and CDKN2D (p19) (**Figure 3C**). This signature is consistent with the transcriptional signature of deep quiescence recently described in fibroblasts in response to altered lysosomal activity in vitro (Fujimaki et al., 2019). Notably, we found a significantly increased ratio of p21 to cyclin D1 in MuSCs from ketone body-injected mice (**Figure S3A**), which is consistent with a higher E2F switching threshold that has previously been shown to be a critical driver of quiescence depth (Kwon et al., 2017). No change was observed in the senescence marker p16 in response to ketone bodies, consistent with the absence of any features of senescence in MuSCs in response to fasting, ketogenic diet, or ketone body injections (**Figure S3B**).

**Figure 3.**
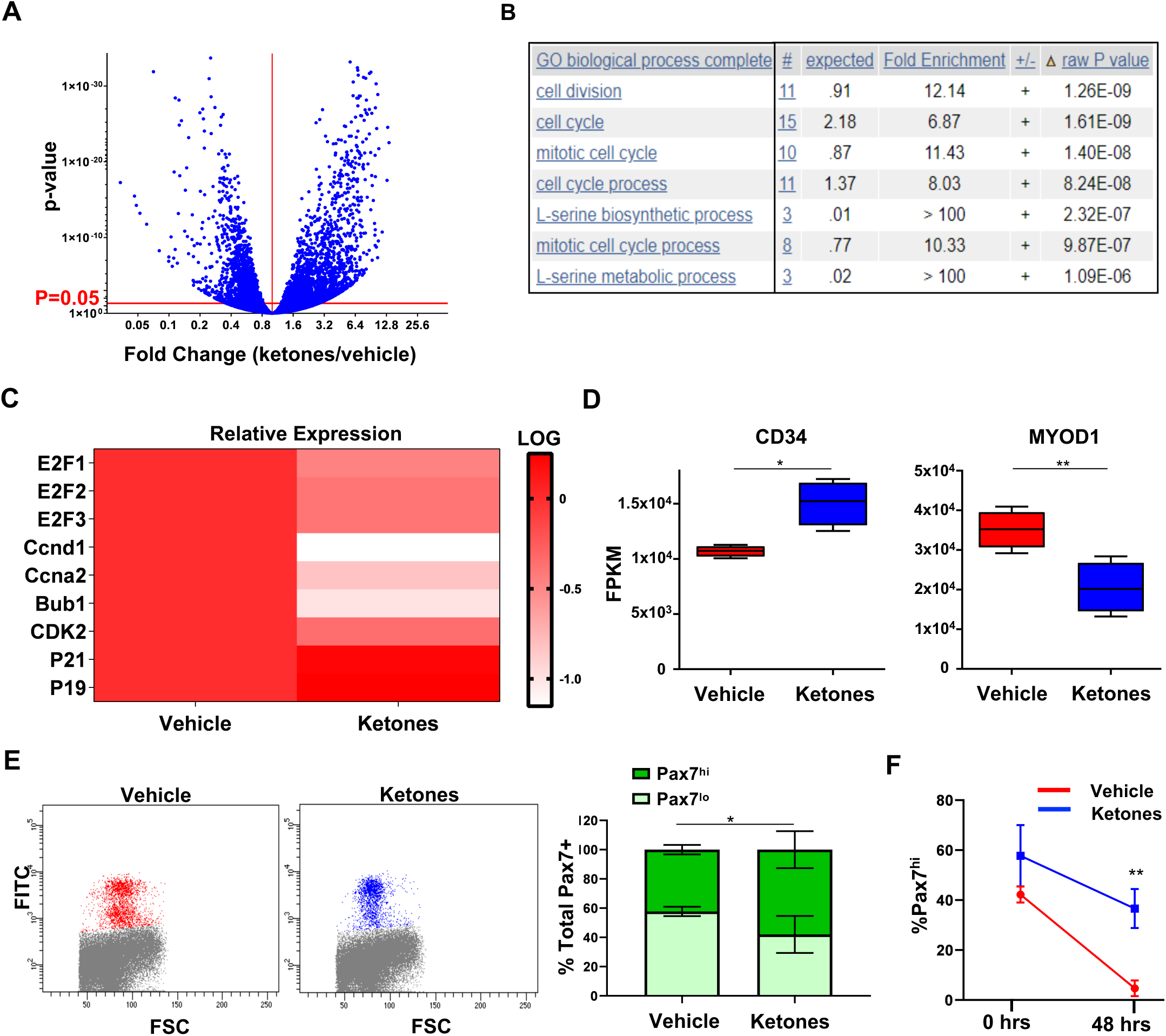
Ketosis promotes a cell cycle and myogenic signature consistent with deep quiescence. **A**) RNA Seq volcano plot comparing differential expression of genes between freshly isolated MuSCs from mice treated with ketone bodies or vehicle. The Y axis is the FDR-normalized p value, and the X axis is the relative fold change of MuSC genes from ketone-treated divided by vehicle-treated mice. Statistically significant changes reside above the red line demarcating a FDR corrected p value of 0.05 (n=4). **B**) Gene ontology analysis comparing enriched biological pathways between freshly isolated MuSCs derived from ketone body- and vehicle-treated mice. **C**) Heat map indicating average relative Log_2_[FPKM] gene expression of select cell cycle genes between freshly isolated MuSCs from ketone body- or vehicle-treated mice (n=4). **D**) Comparative gene expression of CD34 and MYOD1 in freshly isolated MuSCs derived from ketone body- or vehicle-treated mice (n=4). **E**) Representative FACS plots of freshly isolated MuSCs isolated from Tg(Pax7-ZsGreen) mice injected i.p with ketone bodies or vehicle for 1 week (n=3). The Y axis (FITC) shows Pax7 promoter activity and the X axis (FSC) shows cell size. Quantification (right) of the percentage of Pax7^hi^ vs Pax7^lo^ promoter activity in freshly isolated MuSCs from Tg(Pax7-ZsGreen) mice injected with ketone bodies or vehicle (n=3). **F)** Quantification of the percentage of Pax7^hi^ promoter activity in MuSCs isolated from Tg(Pax7-ZsGreen) mice injected with ketone bodies or vehicle. Cells were plated for either 0 or 48 hours at which point Pax7 promoter activity was assessed by FACS. Error bars represent SEM. *p < 0.05; **p < 0.01; ***p < 0.001; ns, not significant. See also Figure S3.

In addition to cell cycle changes, our transcriptomic analysis revealed that freshly isolated MuSCs from ketone-injected mice exhibited lower expression of myogenic genes such as MYF5 and MYOD1(**Figures 3D** **and S3C)**. They also displayed increased expression of CD34, a MuSC stemness marker (Ieronimakis et al., 2010) (**Figure 3D**). Previous work from our lab and others has shown that quiescent MuSCs contain MYOD1 transcripts (de Morree et al., 2017), perhaps allowing quiescent MuSCs to be poised for myogenesis upon activation. Because DQ MuSCs contain less MYOD1 and higher levels of CD34, this may be indicative of MuSCs that are more stem-like and less poised for myogenic commitment. While total Pax7 transcript levels were not significantly altered in the freshly isolated MuSCs from ketone-treated mice **(Figure 3SD)**, we found that freshly isolated DQ MuSCs had a larger percentage of cells with Pax7^hi^ promoter activity (**Figure 3E**). Our work is consistent with previous findings that have found a strong correlation between Pax7 promoter activity and delayed exit from quiescence (Rocheteau et al., 2012). Additionally, we found that Pax7 activity persisted much longer during the course of activation of MuSCs from ketone-treated mice **(Figures 3F, and S3E)**, consistent with a state of DQ.

Previous work has shown that the Pax7^hi^ subpopulation of MuSCs has a higher capacity to seed the MuSC niche than the Pax7^lo^ population (Rocheteau et al., 2012). To assess the engraftment potential of DQ MuSCs compared to control MuSCs, we tested whether DQ cells could outcompete their control counterparts in a competitive transplantation assay. In this paradigm, we co-transplanted distinctly labeled DQ MuSCs and control MuSCs into the same injured recipient muscle. After 28 days, we isolated by FACS those transplanted DQ and control MuSCs that had engrafted. We found that DQ MuSCs consistently outcompeted their control counterparts and were present in higher numbers in the recipient muscle **(Figure S3F)**. These results suggest that the state of MuSC DQ is also characterized by an enhanced capacity for long-term engraftment.

### Ketone-Induced deep quiescence (KIDQ) promotes MuSC resilience

In order to test for the propensity of DQ MuSCs to exit quiescence and enter the cell cycle, we plated MuSCs from ketotic and control mice in culture with the intention of studying in more detail their kinetics of activation. However, surprisingly, we observed that a greater number of MuSCs from ketotic mice were present in culture after 48 hours of activation compared to their control counterparts (**Figure S4A**). Having already observed that DQ MuSCs take longer to break quiescence and enter the cell cycle, we surmised that the most likely explanation for this increase in cell number was not due to faster expansion, but rather an enhanced ability to survive the stress of the activation process (Liu et al., 2018). Consistent with this hypothesis, we found that DQ MuSCs exhibited a dramatic decrease in cell death in culture (**Figures S4B-S4D**), suggesting that the state of DQ is one of a marked increase in resilience. We next asked whether these DQ MuSCs could also resist other forms of stress as well. Indeed, we found that MuSCs isolated from ketone body-injected mice were much more resistant to oxidative stress as well as that of acute nutrient deprivation (**Figures 4A** **and S4E**). Together, these data highlight another key property of DQ cells, and that is one of increased resilience and general stress resistance.

**Figure 4.**
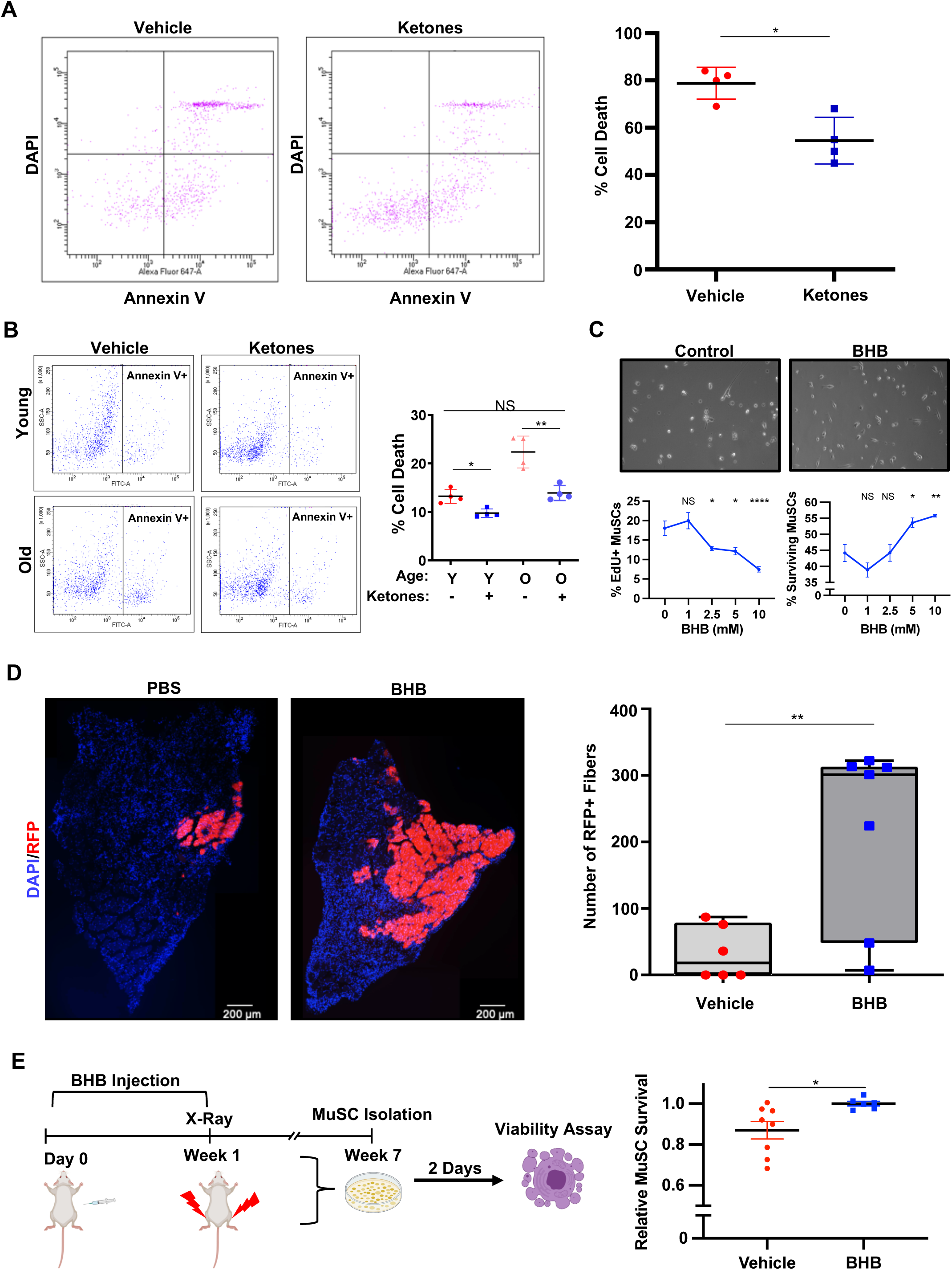
Ketone-Induced deep quiescence (KIDQ) promotes MuSC resilience. **A**) Representative FACS plots (left) of freshly isolated MuSCs from ketone body- or vehicle-treated mice 2 hours following H_2_O_2_ challenge. MuSCs were stained with annexin V (X-axis) and DAPI (Y-axis) to measure cell death (n=4). Quantification (right) of cell death as the percentage of cells that are both propidium iodide and annexin V positive. **B**) Representative FACS plots (left) of MuSCs isolated from young and old mice following one week of in vivo treatment with ketone bodies or vehicle. After isolation, cells were grown for 48 hours in culture and subsequently stained with annexin V to measure cell death (n=4). Quantification (right) of cell death by measuring the percentage of cells that are annexin V positive. **C**) Representative brightfield images (top) of MuSCs following 96 hours of growth in the presence of vehicle control or 10 mM BHB. Quantification of percent EdU incorporation in MuSCs after 48 hours (bottom left; n=4 or 5) and percent MuSC survival after 96 hours (bottom right; n=5) in growth media containing racemic (R/S) BHB at different doses. **D**) Representative fluorescence images (left) of RFP-expressing MuSCs treated in culture with 10 mM BHB or vehicle for 60 hours and subsequently transplanted into pre-injured and pre-irradiated recipient TA muscle (n=6-7 per group). Cryosections harvested 2 weeks after transplantation were analyzed for RFP expression, and the number of RFP positive fibers was quantified (right). **E**) Work flow (left) and quantification (right) of in vivo MuSC resilience in which mice were treated with BHB and then exposed to X-ray irradiation to the hindlimbs. MuSCs from irradiated muscles were isolated and analyzed for viability in culture. (n=6-8 per group). Error bars represent SEM. *p < 0.05; **p < 0.01; ***p < 0.001; ns, not significant. See also Figure S4.

We have recently shown that old MuSCs have an impaired ability to survive the transition between quiescence and activation (Liu et al., 2018). Given the resistance of DQ MuSCs to activation-induced death, we wanted to know if we could rescue the known survival impairments of old MuSCs by promoting KIDQ. In order to address this, we injected a cohort of 24-month-old mice with either vehicle or ketone bodies for one week, at which point we sacrificed the mice and isolated MuSCs from the hindlimbs. After plating the cells for 48 hours in culture, we again measured cell death. Quite strikingly, we found that ketone body treatment could indeed rescue the survival defects that are characteristic of old MuSC activation (**Figure 4B**).

We next wanted to know if KIDQ was a result of the direct effect of the ketone bodies on the MuSCs themselves. Indeed, treatment of isolated MuSCs with BHB (the primary ketone body produced during fasting) produced a striking improvement in cell survival and a concomitant decrease in the rate of S phase entry in a dosage-dependent manner, consistent with the DQ phenotype observed in MuSCs isolated from ketotic mice (**Figure 4C**). Similarly, directly treating isolated human MuSCs with BHB also delayed quiescence exit and reduced cell death in culture (**Figure S4F-S4G**). To determine the functional relevance of this BHB-driven cell-intrinsic survival enhancement to in vivo potency of murine MuSCs, we performed a stem cell transplantation assay. RFP-labeled MuSCs were purified and maintained for 60 hours in medium supplemented with BHB or control medium and then transplanted into injured, irradiated muscles of recipient mice. Fourteen days later, we examined host muscles for the presence of RFP-labeled muscle fibers. Compared to control MuSCs, BHB-treated MuSCs contributed much more robustly to the formation of new muscle fibers (**Figure 4D**). Our finding that ketosis causes reduced expression of genes involved in proliferation and myogenic differentiation (**Figures 3B-3D** and **S3C**) is consistent with the interpretation that increased myofiber formation results from improved MuSC survival during and after transplant, rather than increased proliferation of transplanted cells. To directly test whether BHB treatment leads to enhanced MuSC resilience, we assessed whether ketosis enhances stem cell protection in vivo by exposing vehicle- or BHB-treated mice to genotoxic irradiation. MuSCs from irradiated muscles were then isolated, cultured, and assayed for viability. MuSCs from animals treated with BHB showed significantly reduced death following in vivo x-ray exposure (**Figure 4E**). Taken together, these data suggest that the ketone body BHB produces a highly resilient DQ state in MuSCs.

### KIDQ promotes deep quiescence in MuSCs via an HDAC1-p53 axis

We hypothesized that the effect of ketone bodies in MuSCs was due to changes in the metabolic activity of these quiescent cells. Previous transcriptomic analyses from our lab and others have revealed a host of unique metabolic changes that define the quiescent state (Cheung and Rando, 2013; Fukada et al., 2007). To probe the metabolism of quiescence at the protein level, we employed an untargeted shotgun proteomics analysis of quiescent MuSCs, comparing the data to those from activated MuSCs. Screening for quiescence-enriched metabolic enzymes, we found that OXCT1, the rate limiting enzyme in ketone body metabolism (Catabolism, 2008), was highly expressed in the quiescent state (**Figures 5A** **and S5A**). This suggested to us that ketone bodies may be promoting DQ in MuSCs via their conversion to acetyl CoA. To address this, we generated a conditional OXCT1 KO mouse (*OXCT1*^fl/fl^;*Pax7*^CreER/+^;*ROSA26*^eYFP/+^ ) to specifically ablate OXCT1 in MuSCs following tamoxifen injection (**Figures S5B and S5C**) (Cotter et al., 2013; Nishijo et al., 2009). Two weeks after the completion of tamoxifen dosing, we injected the control and OXCT1^cKO^ mice with ketone bodies 3 times per day for 7 days. To our surprise, we found that there was no significant difference in the induction of DQ in MuSCs in response to exogenous ketone body treatment between the two strains (**Figures 5B** **and S5D-S5F**). This finding suggested to us that ketone bodies are likely acting non-metabolically to elicit DQ in these cells.

**Figure 5.**
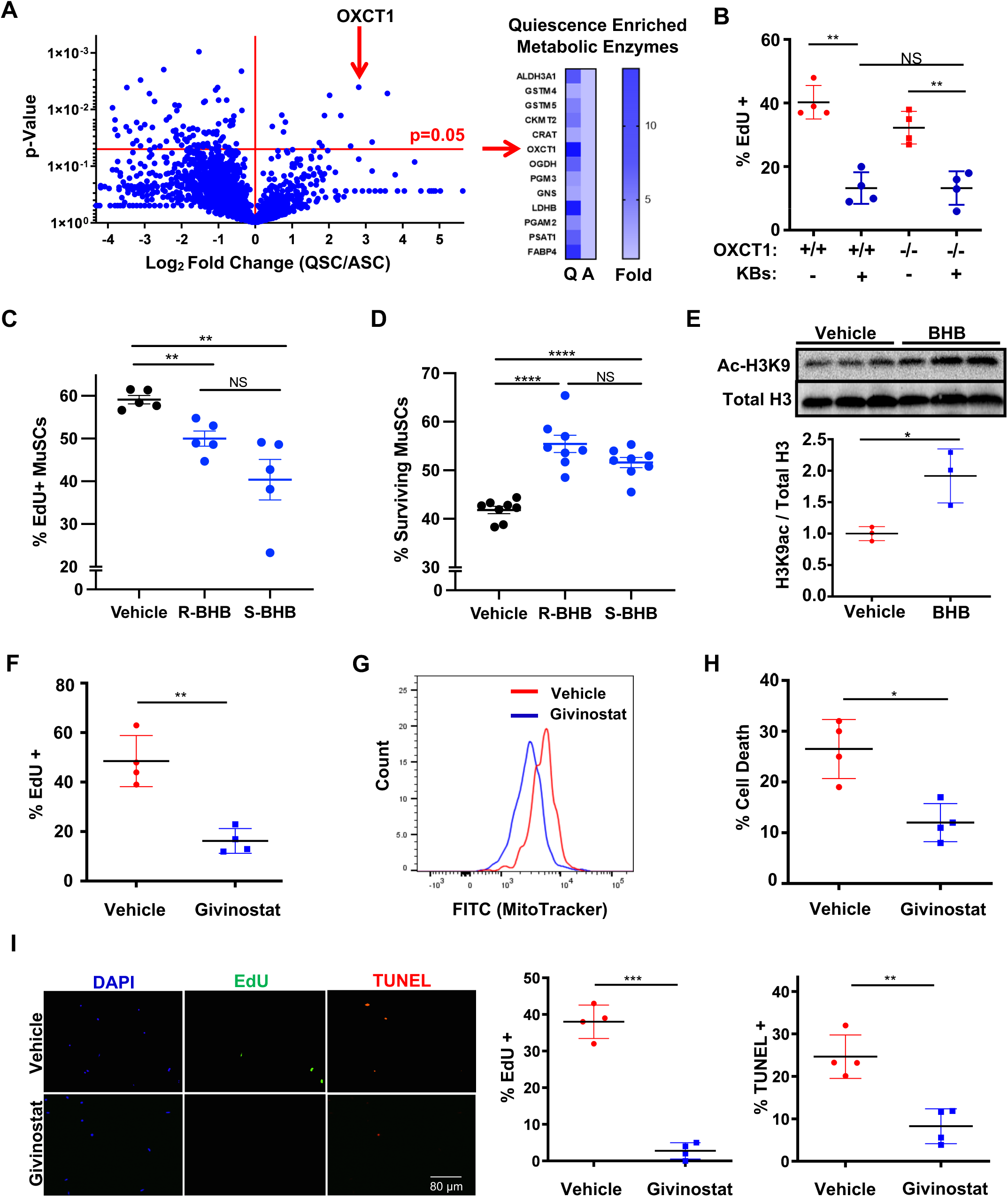
Ketone bodies elicit DQ via a nonmetabolic HDAC inhibitory mechanism. **A**) Shotgun proteomics volcano plot (left) comparing relative spectral count values between quiescent and activated MuSCs. The Y axis represents the p value and the X axis represents the ratio of protein spectral counts of quiescent divided by activated MuSCs. Statistically significant protein changes reside above the red line demarcating a p value of 0.05. OXCT1 is identified and is shown in a heat map (right) alongside other quiescence-enriched metabolic enzymes. Quiescence-enriched proteins in the heat map are shown as relative fold change normalized to activated MuSCs (n=2; 20 pooled mice per replicate). **B**) Quantification of EdU incorporation in MuSCs derived from OXCT1 cKO or WT mice that had been treated with either ketone bodies or vehicle for 1 week. Cells were maintained in culture for 48 hours in the continuous presence of EdU (n=4). **C**) Percent EdU incorporation of MuSCs after 48 hours in culture with growth media that contained vehicle or 2.5 mM of the indicated stereoisomer of BHB. **D**) Survival rates of MuSCs after 96 hours in culture with growth media that contained vehicle or 5 mM of the indicated stereoisomer of BHB. **E**) Representative western blot analysis (left) of histone 3 acetylation at lysine 9 in MuSCs in response to 10 mM BHB treatment for 24 hours in culture. Total histone H3 is used to control for protein loading (n=3 per group). Quantification (right) of relative histone H3 acetylation at lysine 9 normalized first to total H3 and then to vehicle control. **F**) Quantification of EdU incorporation in MuSCs isolated from mice treated with givinostat (10 mg/kg i.p. daily) or vehicle for 1 week. Following isolation, MuSCs were maintained in culture for 48 hours in the continuous presence of EdU (n=4). **G**) Representative FACS plot of MitoTracker Green intensity in freshly isolated MuSCs from givinostat- or vehicle-treated mice (n=4). **H**) Quantification of cell death in MuSCs from givinostat- or vehicle-treated mice. Cells were grown in culture for 48 hours and subsequently stained for annexin V and propidium iodide. Cell death was quantified by the percentage of cells that were positive for annexin V and propidium iodide. **I**) Representative immunofluorescence images (left) of MuSCs treated with givinostat or vehicle in culture for 48 hours. Cells were grown in the continuous presence of EdU, and after 48 hours were stained for EdU (488 channel) and TUNEL (594 channel). Nuclei were stained with DAPI. Quantification of EdU incorporation (middle) and cell death (right). Error bars represent SEM. *p < 0.05; **p < 0.01; ***p < 0.001; ns, not significant. See also Figure S5.

Since non-metabolic signaling functions have been attributed to both AcAc and BHB (Juge et al., 2010; Newman and Verdin, 2014; Shimazu et al., 2013), we next wanted to distinguish which of the two metabolites is the primary driver of DQ during ketosis. Addressing this question by simply treating MuSCs with BHB and AcAc separately is confounded by the fact that the two molecules readily interconvert through the enzymatic activity of β-hydroxybutyrate dehydrogenase (BDH1). However, BDH1 is known to specifically convert the endogenously produced R enantiomer of BHB into AcAc but does not metabolize the S-BHB (Lehninger et al., 1960). We took advantage of this limitation in the enzymatic activity of BDH1 by treating MuSCs with the non-metabolizable S-enantiomer to determine whether BHB could promote DQ absent its conversion into AcAc. Indeed, we found that treatment of MuSCs with S-BHB was sufficient to delay quiescence exit, as measured by EdU incorporation (**Figure 5C**) and to increase survival rate of cultured MuSCs by >20% (**Figure 5D**). These data support our previous finding that ketosis promotes MuSC DQ independent of ketone body metabolism by OXCT1 and establish that BHB alone is sufficient to promote MuSC DQ, absent its conversion to AcAc.

The best characterized non-metabolic function of BHB is as an endogenous inhibitor of histone deacetylases (HDACs) (Cheng et al., 2019; Newman and Verdin, 2014; Shimazu et al., 2013). To probe whether or not BHB might be acting as an HDAC inhibitor in MuSCs, we treated isolated MuSCs with BHB and measured acetylation of histone H3 (a well characterized HDAC substrate) at lysine 9. We found that BHB administration resulted in significant elevations in H3K9 acetylation, consistent with its role as a potential HDAC inhibitor (**Figure 5E**). Similarly, fasting-induced ketosis resulted in elevated H3K9 as well as H3K14 acetylation (**Figures S5G-S5H**). If the effects of BHB on promoting MuSC DQ were indeed the result of HDAC inhibition, we surmised that HDAC inhibition might phenocopy the effects of fasting, the ketogenic diet, and exogenous ketone body administration on the promotion of DQ in MuSCs. To address this, we injected mice intraperitoneally with an HDAC inhibitor in clinical use, givinostat, or with a vehicle control for one week. We found that MuSCs isolated from mice injected with givinostat displayed all of the DQ hallmarks seen in MuSCs from ketotic mice (**Figures 5F, 5G, and S5I-S5J**). Givinostat treatment had no effect on the body weight or ketone levels of the mice (**Figures S5K and S5L**). Consistent with the hallmarks of the DQ phenotype, MuSCs isolated from givinostat-injected mice exhibited a remarkable ability to survive in culture and to resist the stress of acute nutrient deprivation compared to control MuSCs (**Figures 5H, S5M, and S5N**). Notably, as we observed with fasting, the effects of givinostat on MuSCs perdured multiple days after treatment was discontinued (**Figures S5O-S5R**). To examine whether or not this phenotype was the result of a direct effect of givinostat on the MuSCs themselves, we isolated MuSCs and treated them in culture with givinostat for 48 hours. Similar to what we observed with ex vivo BHB treatment, we found that treatment of isolated MuSCs with givinostat ex vivo resulted in a dramatic enhancement of MuSC survival along with a substantial delay in the entry into S phase (**Figure 5I**).

In order to narrow down exactly which class of HDACs, when inhibited, is sufficient to promote MuSC DQ, we performed an HDAC inhibitor screen using a chemical library of well-characterized HDAC inhibitors. Our screen revealed that HDAC class 1 (and very likely HDAC1 within class 1) was highly enriched among inhibitor targets that were able to promote DQ **(Figures S5S-S5V)**. To confirm the findings of our screen in vivo, we injected mice with inhibitors from the screen that spanned the different classes of HDAC inhibitors. Consistent with the findings from our screen, we found that only HDAC class I inhibitors, when injected intraperitoneally, were able to elicit the phenotypic hallmarks of DQ in MuSCs **(Figures S5W and S5X)**.

We next wanted to determine exactly which of the HDAC1 targets, when hyperacetylated in response to HDAC inhibition, might be driving the DQ phenotype in MuSCs. To address this question, we performed a pan acetyl-lysine western blot to assess for any pattern of increased protein acetylation in response to fasting or ketone body treatment. Intriguingly given that p53 is a canonical HDAC1 target (Ito et al., 2002), we found a protein of approximately 53 kD that was most prominently hyperacetylated in response to ketosis (**Figure 6A**). Mouse p53 can be acetylated at various residues, including K379 (K382 in human) (Sakaguchi et al., 1998), and K379 acetylation is known to promote DNA binding activity of p53, induce apoptosis in cancer cells, and promote survival in certain non-cancer cells (Brochier et al., 2013; Sakaguchi et al., 1998). Using an antibody specific to acetyl-p53 K379, we confirmed hyperacetylation of p53 by both western blot of FACS-purified MuSCs and immunofluorescence analysis of MuSCs on isolated myofibers in response to fasting, the ketogenic diet, and ketone body injections **(Figure 6B-6C, and S6A-S6B)**. These findings are consistent with recent work suggesting that liver p53 is also hyperacetylated in response to both short-term fasting and ketogenic diet treatment (Roberts et al., 2017). In addition, we identified a robust p53 transcriptional signature in MuSCs by gene set enrichment analysis in response to ketosis (**Figure 6D**). Recent work from our lab has highlighted the unique and unexpected role of p53 in promoting the survival of MuSCs (Liu et al., 2018). We therefore surmised that acetylation and activation of p53 via HDAC1 inhibition might be one of the pathways contributing to the DQ state of enhanced MuSC resilience in response to ketosis and givinostat treatment.

**Figure 6:**
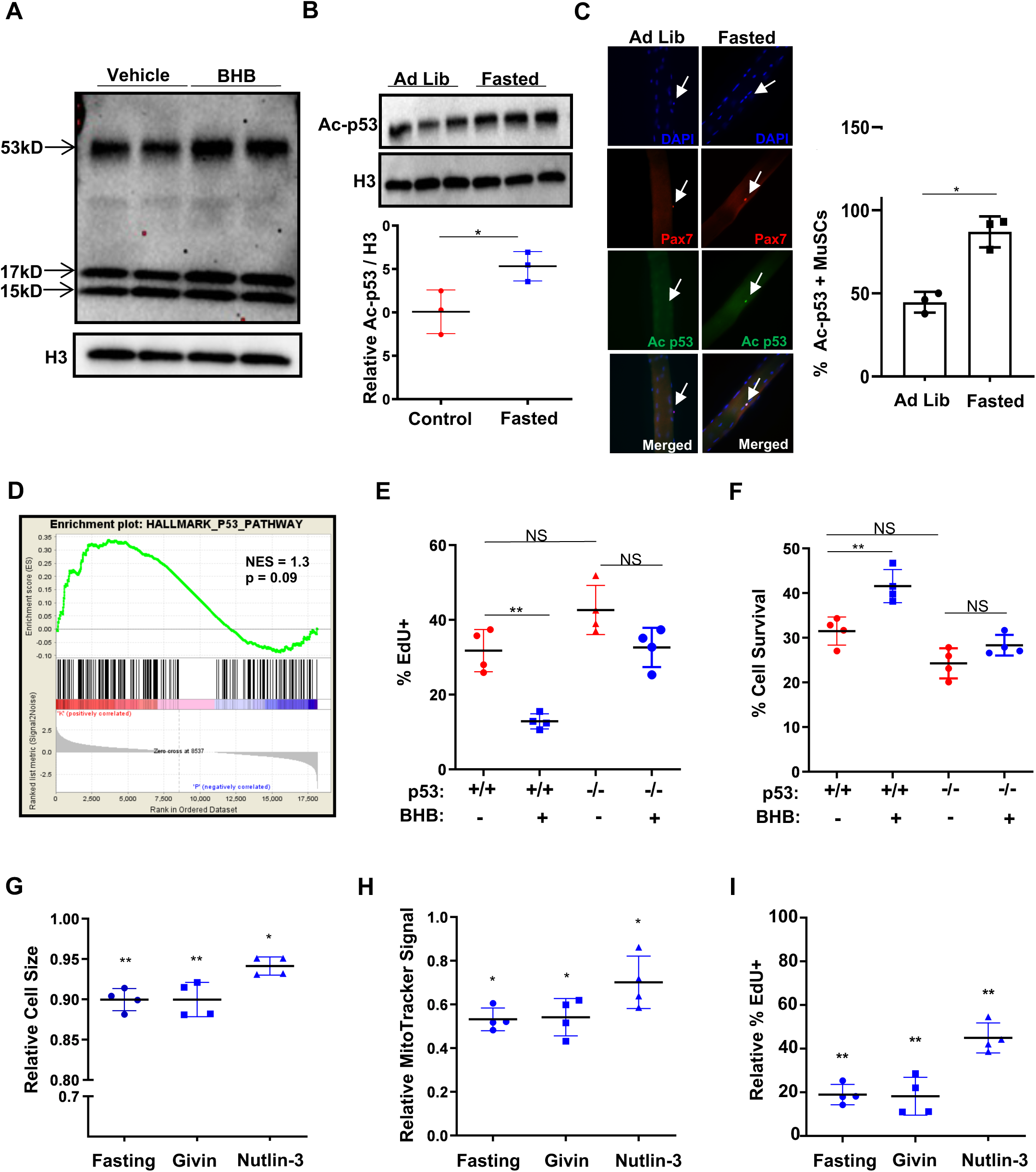
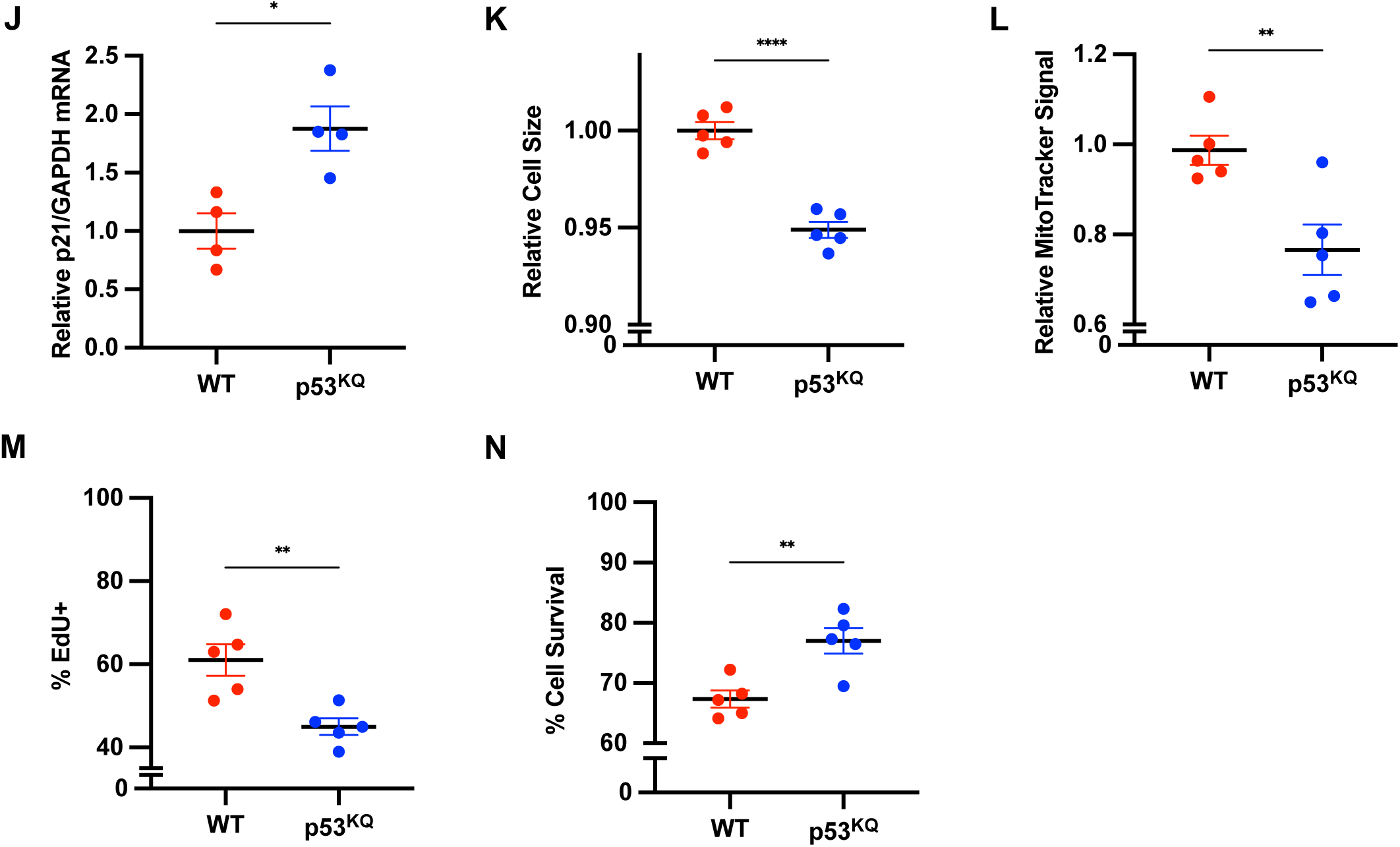
p53 is both necessary and sufficient to promote MuSC DQ. **A**) Representative western blot showing pan acetyl lysine levels in freshly isolated MuSCs from mice treated with BHB or vehicle for one day. Bands at 53 kD, 17 kD, and 15 kD were identified as exhibiting increased acetylation upon in vivo BHB treatment. H3 was used as a control for protein loading. **B**) Representative western blot (top) showing acetyl-p53 levels in MuSCs isolated from fasted or ad lib fed mice. Total H3 protein was used as a loading control. Quantification (bottom) of relative acetyl-p53 levels normalized first to total H3 and then to ad lib fed controls (n=3). **C**) Immunofluorescence staining (left) of Pax7 and acetyl-p53 on single isolated myofibers isolated from fasted and ad lib fed control mice. Nuclei were stained with DAPI. Arrows indicate MuSCs. Quantification (right) of percent of Pax7 positive MuSCs that stained positive for acetyl-p53. **D**) Enrichment plot for the p53 pathway gene set. NES stands for normalized enrichment score. **E**) Quantification of EdU incorporation in MuSCs derived from p53 cKO or WT mice treated with BHB in culture. Cells were treated with either 10 mM BHB or vehicle for 48 hours in the continuous presence of EdU (n=4). **F**) Quantification of cell survival in MuSCs isolated from p53 cKO or WT mice treated with BHB in culture. Cells were treated with either 10 mM BHB or vehicle for 96 hours and subsequently stained for annexin V and PI. Percent cell survival was quantified as the percentage of cells that stained negative for both annexin V and PI. **G-I**) Quantification of (G) relative cell size (based on forward scatter from FACS) in freshly isolated MuSCs, (H) relative MitoTracker intensity in freshly isolated MuSCs, and (I) percent EdU incorporation at 48 hours in MuSCs derived from mice that were fasted for 60 hours or injected with givinostat (10 mg/kg daily) or with the p53 activator nutlin-3 (20 mg/kg daily) for 1 week. Values from each treatment condition were normalized to respective ad lib or vehicle controls (n=4). **J**) Quantification of p21 transcript levels by qPCR of mRNA extracted from MuSCs of WT and p53^KQ^ mutant mice. p21 transcript levels were normalized to GAPDH transcript levels (n=4). **K-N**) Quantification of (K) relative cell size (forward scatter), (L) MitoTracker Signal, (M) percent EdU incorporation, and (**N**) percent survival of MuSCs from WT and p53^KQ^ mutant mice (n=5). Error bars represent SEM. *p < 0.05; **p < 0.01; ***p < 0.001; ****p < 0.0001; ns, not significant. See also Figure S6.

To test for a potential role of p53 in mediating any of the phenotypes of KIDQ, we used a Pax7CreER driver to specifically ablate p53 in MuSCs as previously described (Liu et al., 2018; Nishijo et al., 2009). Two weeks after the completion of tamoxifen dosing, MuSC-specific p53 KO mice and WT controls were fasted or ad libitum fed for 60 hours, at which point their MuSCs were isolated for analysis. We found that key hallmarks of DQ, including reduced cell size, decreased mitochondrial content, and delayed S phase entry, were partially abrogated in response to the ablation of p53 in MuSCs **(Figures S6C-S6G)**. In order to determine if p53 were mediating the direct effects of BHB on MuSCs, we isolated p53-ablated MuSCs and WT controls, and we treated the cells ex vivo with BHB. We found that that p53 KO MuSCs were largely insensitive to BHB-induced changes in cell cycle entry and survival (**Figures 6E** **and 6F)**

To test for the sufficiency of p53 activity to promote DQ, we injected mice intraperitoneally with vehicle or with nutlin-3, a p53 activator (Vassilev et al., 2004). We found that activating p53 in vivo could indeed recapitulate the DQ phenotypes of MuSCs (**Figures 6G-6I and S6H-S6I**) that we had seen previously with both ketosis as well as HDAC inhibition. Additionally, consistent with what we found with BHB and givinostat treatments, treating isolated MuSCs with nutlin-3 for 48 hours delayed cell quiescence exit and promoted resilience in cultured MuSCs (**Figures S6J and S6K**). Activity of p53 can be regulated through post-translational acetylation of various lysine residues in the C-terminus of the protein, including lysine 379 (382 in humans) (Brochier et al., 2013). Recent studies have found that the stabilizing effects of acetylation can be mimicked by mutating various C-terminal lysine residues to glutamine (p53^KQ^) (Wang et al., 2016). To directly demonstrate the ability of p53 acetylation to mimic the effects of ketosis, we tested whether MuSCs isolated from these p53^KQ^ mutant mice exhibit the hallmarks of DQ. Indeed, p53^KQ^ MuSCs present evidence of elevated p53 activity as well as reduced cell size and mitochondrial content at time of isolation, delayed S phase entry, and enhanced survival (**Figures 6J-N**). Collectively, our results highlight both the necessity and sufficiency of p53 activity in promoting MuSC DQ.

## DISCUSSION

Fasting elicits a host of beneficial effects on multiple cell and tissue types (Brandhorst et al., 2015), including functional improvements in certain adult stem cell populations. Here we show that fasting causes MuSCs to enter a perdurant, highly protective deep quiescent state in which they are resistant to various forms of extreme stress and display improved self-renewal. These results are consistent with previous findings that cycles of periodic fasting increase HSC self-renewal and protects HSCs against damage from chemotherapy reagents (Cheng et al., 2014). Although there have been conflicting reports regarding the effect of caloric restriction on muscle regeneration, (Boldrin et al., 2017; Cerletti et al., 2012), we have definitively shown that fasting induces MuSC DQ, a state characterized by delayed cell cycle entry and delayed expression of the myogenic program, resulting in delayed in vivo myogenesis after injury. Intriguingly, these findings shed light on a dual nature of fasting in the context of MuSC function. Whereas fasting appears detrimental to the rate of muscle regeneration following acute injury, fasting may improve long term tissue maintenance by enhancing the resilience and self-renewal of MuSCs via the induction of a deep quiescent state. This duality between the benefits and drawbacks of dietary restriction is also supported by recent studies in the immunology field. It was shown that caloric restriction results in T cell homing to the bone marrow and improved clearance of infection (Collins et al., 2019), and fasting decreases the circulating inflammatory monocyte population, reducing inflammation (Jordan et al., 2019). However, fasting also results in death of germinal center B cells and perturbs the antigen-specific IgA response (Nagai et al., 2019). Our work is consistent with recent work from our lab and others which suggests that long-term regenerative capacity of muscle tissue depends heavily on maintenance of the quiescent state to prevent spontaneous activation and depletion of the MuSC pool through cell death or precocious differentiation (Bjornson et al., 2012; Conboy et al., 2003; Liu et al., 2018; Mourikis et al., 2012). In this context, dietary interventions such as fasting or the ketogenic diet may help maintain muscle regenerative capacity throughout organismal lifespan. Recent studies that explored the anti-aging effects of dietary ketosis in mice found extended lifespan and preserved muscle mass after long-term consumption of a ketogenic diet (Roberts et al., 2017; Wallace et al., 2021). While DQ could potentially prevent MuSC decline during aging via improved resilience, MuSC maintenance is unlikely to contribute to this reduction in sarcopenia in the absence of injury (Fry et al., 2015). The ability of ketogenic diets to prevent age-related muscle atrophy might be better explained by direct effects of diet on promoting resilience of mature muscle, independent of effects on MuSCs.

Consistent with previous work from our lab and others (Fujimaki et al., 2019; Kwon et al., 2017; Rodgers et al., 2014), the data presented here suggest a considerable degree of plasticity in the quiescent state. Furthermore, our data reveal that perhaps a central feature of quiescence plasticity is a balance between the readiness to activate and cellular resilience. In support of this model, previous work from our lab has reported on a separate state of quiescence (G_Alert_), induced by distant tissue injury, that is defined primarily by a state of shallow quiescence and therefore an enhanced readiness to activate and contribute to myogenesis (Rodgers et al., 2014). We have recently shown that MuSCs in G_Alert_ suffer from a defect in resilience manifested as a failure of long term maintenance (Haller et al., 2017). Whereas recent work has uncovered the existence of DQ in other cell types (Fujimaki et al., 2019; Kwon et al., 2017), including a deeply quiescent subpopulation of HSCs (Matsuoka et al., 2011), our work has highlighted a novel means by which dietary intervention can promote DQ in adult stem cells. Moreover, our work indicates that quiescence depth may be inextricably linked to cellular stress resistance and that changing the depth of quiescence may change the ability of a MuSC to respond to stress. Consistent with this positive correlation between depth of quiescence and resilience, a recent study found that enhancing quiescence in HSCs by genetic deletion of the transcriptional regulator ELF4 promotes cellular resistance to chemotherapy- and radiation-induced damage (Lacorazza et al., 2006).

In addition to the hormonal and metabolic changes that accompany nutrient deprivation, fasting is characterized by a dramatic elevation in blood ketone levels (Han et al., 2018). Here we show that the administration of ketone bodies themselves, absent any other feature of nutrient deprivation, is sufficient to elicit DQ of MuSCs. Evolutionarily, elevated ketone bodies in the blood may be a rapid way to signal to cell types throughout the body to allocate resources to survival mechanisms, like increased resilience, during a time of nutrient deprivation. Consistent with our findings in MuSCs, this role of exogenous ketone bodies in promoting a quiescent state was recently elucidated in vascular cells (Han et al., 2018). The authors showed that BHB was sufficient to promote quiescence while simultaneously preventing senescence of these cells. Other studies have similarly shown that many of the beneficial effects of nutrient deprivation in other cell and tissue types can also be attributed, in large part, to the accompanying ketosis (Cheng et al., 2019; Huang et al., 2018; Veech et al., 2017). For example, a recent study showed that the neuroprotective and lifespan-extending properties of CR are in part mediated by the ketone bodies themselves (Huang et al., 2018). Interestingly, however, one recent study has reported that a localized intramuscular injection of acetoacetate during muscle regeneration can promote MuSC proliferation and accelerate muscle regeneration through activation of the MEK1-ERK1/2-cyclin D1 pathway (Zou et al., 2016). Notably, this study focused only on the effects of the acetoacetate immediately following injury, when MuSCs are proliferating, without exploring its role during quiescence. Additionally, this study did not look at the effects of systemic ketosis or its predominant ketone body, BHB, on MuSCs. As such, this study does not address the effects of ketosis on quiescent stem cell function.

Although the canonical function of ketone bodies is as a metabolic fuel source and precursor for the production of mitochondrial acetyl CoA, our work indicates that the role of ketone bodies in promoting DQ within MuSCs is not a result of their metabolism to acetyl CoA. Indeed, consistent with findings in other cell and tissue types (Cheng et al., 2019; Newman and Verdin, 2014; Shimazu et al., 2013; Yang et al., 2014), our data suggest that ketone bodies, and specifically the ketone BHB, may act via inhibiting HDAC activity. Specifically, our data indicate that class I HDAC inhibition, and specifically HDAC1 within class I, may be driving the DQ of these cells. A number of previous studies have demonstrated the role of pharmacological HDAC inhibition in controlling stem cell fate and function (Cheng et al., 2019; Leibowitz et al., 2018). Here we present evidence in support of the notion that reduction of HDAC activity by an endogenous inhibitor, namely circulating BHB, may underlie the effects of fasting on a stem cell population.

Our work has elucidated the role of one particular HDAC1 substrate, p53, in promoting MuSC DQ. Whereas multiple previous studies have shown that p53 activation may be coincident with fasting and ketosis (Prokesch et al., 2017; Roberts et al., 2017), none of these studies have elucidated a clear functional role for p53 in this context. We have shown here that p53 activation is both necessary and sufficient for MuSCs to enter the state of DQ. In particular, we have shown that p53 activation is sufficient to promote the enhanced survival of MuSCs. This finding on the protective role of p53 is consistent with previous work from our lab and others elucidating the seemingly paradoxical role of p53 and p53 family members in promoting the survival of multiple different stem and progenitor cell types including neural stem cells and intestinal progenitor cells (Agostini et al., 2010; Liu et al., 2018, 2009). Additionally, our work linking p53 with cellular quiescence in MuSCs is consistent with established role of p53 in mediating the enhanced quiescence of HSCs as well as promoting quiescence in other proliferating cell types via the up-regulation p21, a p53 target protein (Itahana et al., 2002; McConnell et al., 2016; Morales et al., 2016). The fact that p53 is a mediator of MuSC DQ does not preclude the possibility of other HDAC targets also contributing to this process. Our transcriptional data suggest that ketosis induces the activation of other pathways that are regulated by HDAC1 and that have been implicated in MuSC cellular quiescence, including the mTOR and the Rb/E2F pathways (Luo et al., 1998; Morales et al., 2016; Rodgers et al., 2014). It will be of considerable interest to decipher whether additional molecular and cellular pathways may be working in parallel or synergistically with the p53 pathway to promote DQ in MuSCs.

Recent work has demonstrated that p53 can be post-translationally regulated through β-hydroxybutyrylation (BHBylation), which attenuates acetylation and activation of p53 (Liu et al., 2019). In that study, global p53 acetylation levels were decreased after BHB treatment, but p53 acetylation at individual residues was not determined. This is a notable caveat, as p53 K379 acetylation has a primary role in promoting the anti-apoptotic functions of p53, despite the acetylation status of other residues (Brochier et al., 2013). In our hands, p53 becomes hyperacetylated at K379 during ketosis, indicating that ketosis could conceivably promote acetylation at certain prominent residues to drive p53 activity while global p53 acetylation is downregulated through BHBylation. Consistently, our data show a robust transcriptional signature of upregulated p53 activity after ketone body treatment, suggesting that site-specific acetylation of p53 is perhaps more consequential for regulating p53 transcriptional activity in MuSCs than BHBylation. Furthermore, important differences have been observed in the regulation and activity of p53 between healthy cells and tumor cells (Brochier et al., 2013; Liu and Kulesz-Martin, 2000). This is another notable caveat, as regulation of p53 through BHBylation was studied in large part using U2OS, HCT116, and H1299 cancer cell lines (Liu et al., 2019). While the referenced study did find p53 BHBylation and target gene (PUMA and p21) downregulation in mouse thymus after fasting, p53 hyperacetylation and elevated target gene (DDIT4) expression has been previously observed in liver of mice that consumed a ketogenic diet (Roberts et al., 2017), suggesting that p53 regulation during ketosis is tissue dependent. Future studies could elucidate the influence of BHBylation on site-specific p53 acetylation, how BHBylation and acetylation at different sites might be integrated to modulate p53 transcriptional activity, and how p53 acetylation and BHBylation are differentially regulated in various tissues during ketosis.

Our work indicates that fasting, and ketosis in general, elicits a highly protective state (DQ) in MuSCs that is characterized, among other features, by enhanced resilience. Enhanced stress resistance during KIDQ significantly increases the utility of MuSCs in the context of transplantation. This is consistent with the known benefits of both quiescence and cellular preconditioning for stem cell transplantation (Haider and Ashraf, 2008; Quarta et al., 2016). MuSCs are able to form muscle and self-renew upon transplantation (Sacco et al., 2008), characteristics that are promising for cell therapy for treating muscular disorders such as Duchenne muscular dystrophy and volumetric muscle loss (Quarta et al., 2017; Sun et al., 2020). However, the clinical efficacy of MuSCs is limited because of the low number that can be isolated from human muscle biopsies (Sun et al., 2020). Isolated MuSCs can be expanded in culture, but cultured MuSCs tend to have low engraftment efficiency (Quarta et al., 2016), due at least in part to extensive cell death after transplantation (Skuk and Tremblay, 2003; Sun et al., 2020). We show here that survival of cultured MuSCs can be significantly improved by treatment with BHB, resulting in improved engraftment upon transplantation. Furthermore, we show that self-renewal of transplanted MuSCs can be increased by enhancing the depth of quiescence prior to transplantation. Importantly, in further support of the translational applicability of our work, we have also shown that the effects of BHB on MuSCs can be extended to isolated human MuSCs as well. These studies thus identify an intracellular pathway that can be targeted to improve the utility of MuSCs for transplantation-based therapies.

## KEY RESOURCES TABLE

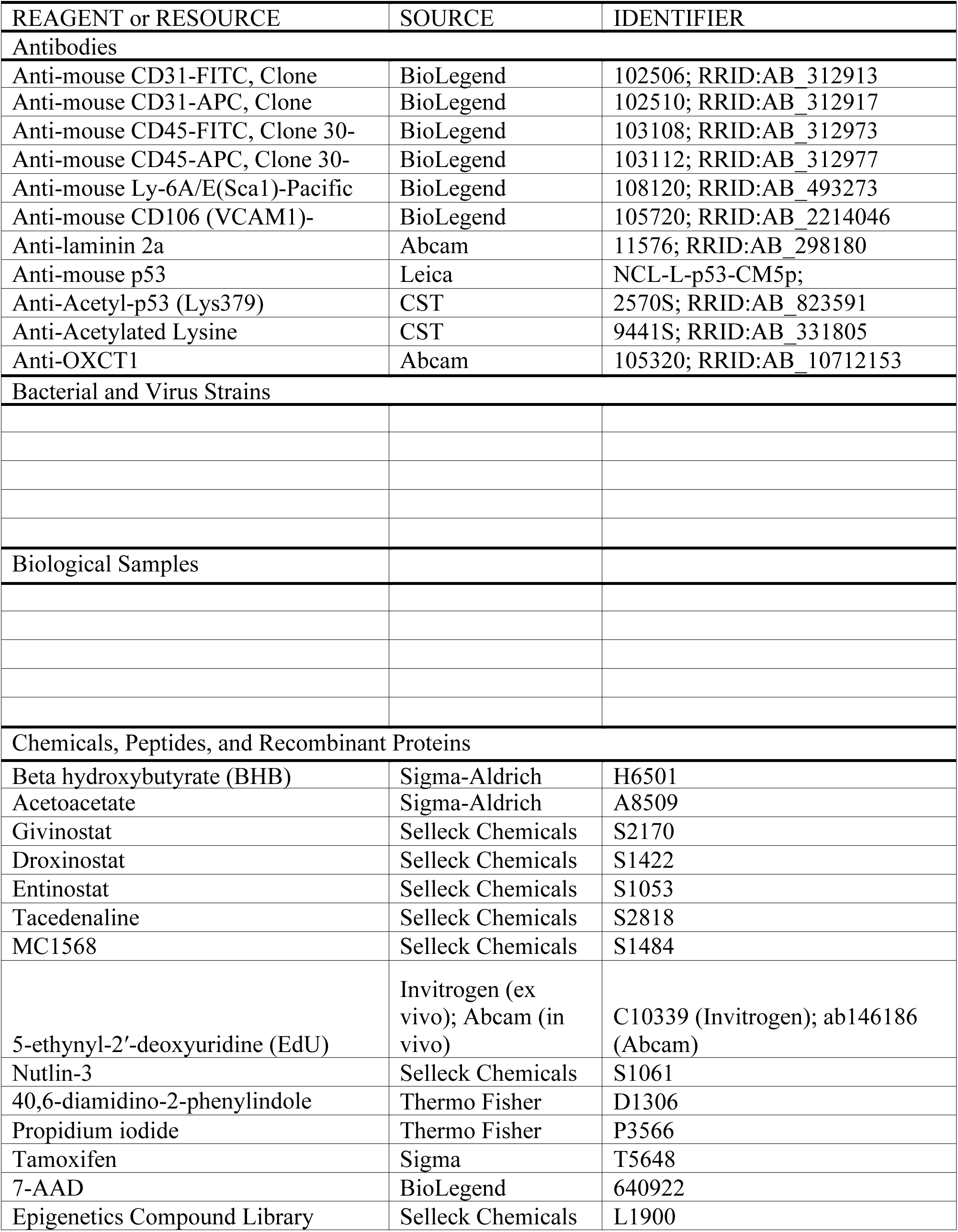
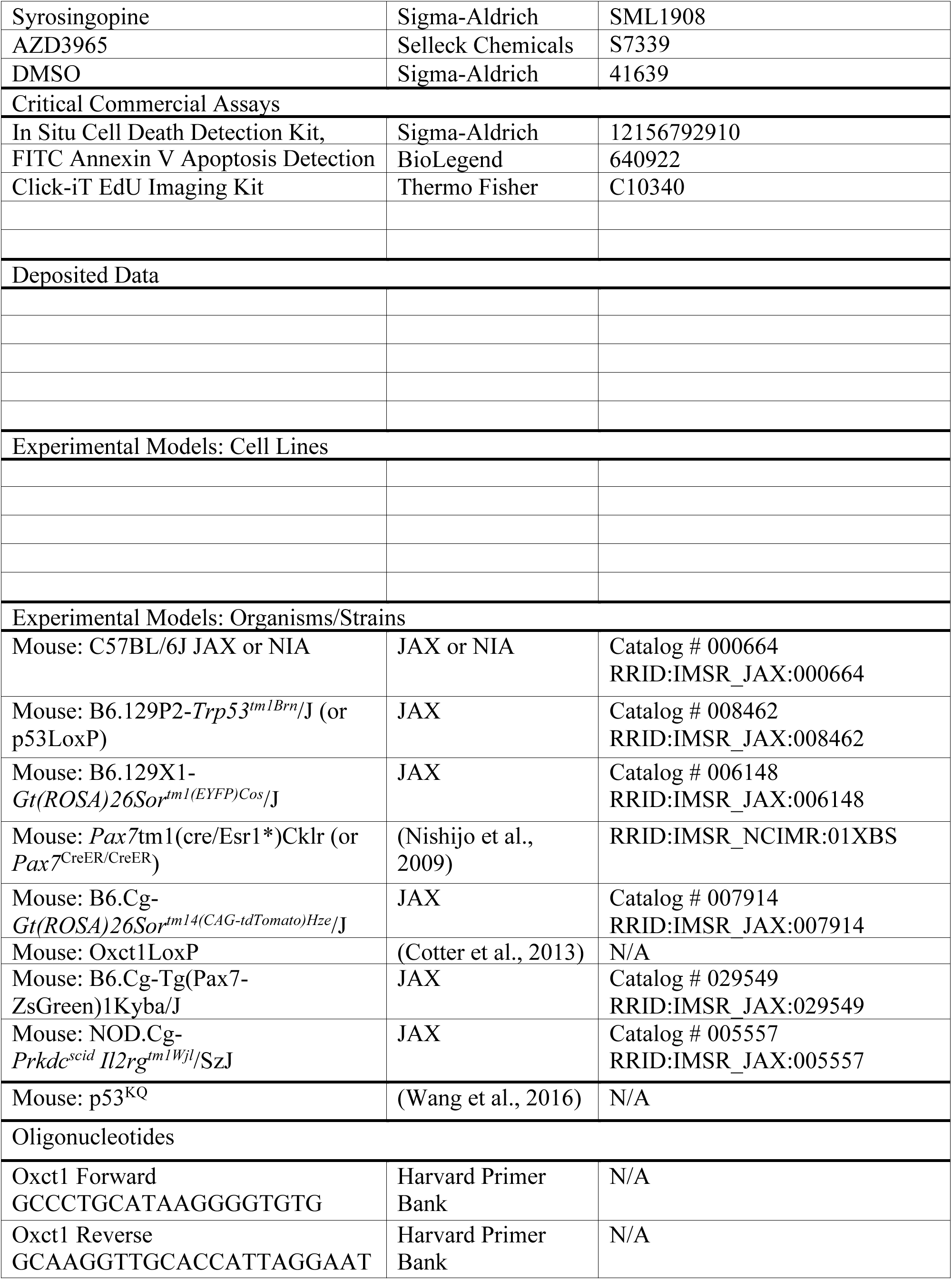
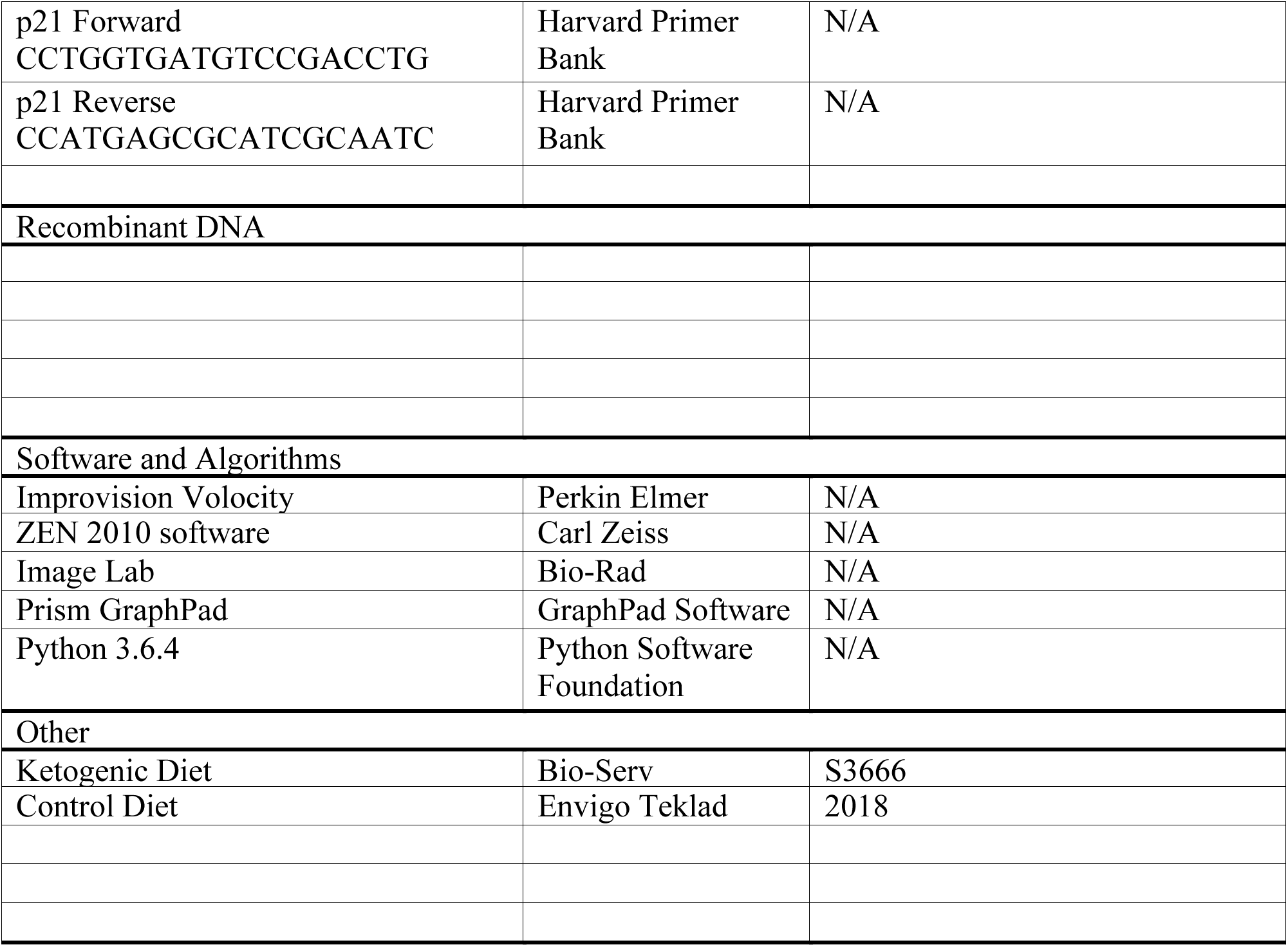

## EXPERIMENTAL MODEL AND SUBJECT DETAILS

### Animals

Animals were housed and maintained in the Veterinary Medical Unit at the Veterans Affairs Palo Alto Health Care System. Animal protocols were approved by the Institutional Animal Care and Use Committee. Young mice used in these studies were 2-4 months old; old mice were 20-24 months old. Young C57BL6 mice were purchased from The Jackson Laboratory (000664). Old C57BL6 mice were obtained from Charles River Laboratories through the National Institute on Aging. NSG mice were purchased from The Jackson Laboratory (005557). *Pax7*^CreER/+^; *ROSA26*^YFP/+^ and *Pax7*^CreER/+^; *ROSA26*^RFP/+^ mice were obtained by crossbreeding *Pax7*^CreER^ mice on the C57BL/6 x 129/SvJ background (Dr. Charles Keller at Oregon Health & Science University) with either *ROSA26*^YFP^ (006148) or *ROSA26*^tdTomato^ (007914) mice purchased from The Jackson Laboratory. *OXCT1*^fl/fl^ mice used to generate MuSC-specific OXCT1 KO mice (*OXCT1*^fl/fl^; *Pax7*^CreER/+^; *ROSA26*^eYFP/+^) were provided by Peter Crawford at Washington University, St. Louis. The *p53*^fl/fl^ mice used to generate MuSC-specific p53 KO mice (*p53*^fl/fl^; *Pax7*^CreER/+^; *ROSA26*^eYFP/+^) were purchased from The Jackson Laboratory (008462). p53^KQ^ mutant mice were provided by Dr. Wei Gu at Columbia University. Mice that were purchased and used in experiments that did not involve in-house husbandry were male. Both male and female mice were bred in-house for experiments involving lineage tracing, genetic recombination, or p53^KQ^ expression.

### Human Skeletal Muscle Specimen

The human muscle biopsy specimen was taken from a 74-year-old Caucasian male. The muscle biopsy sample was taken after patient gave informed consent, as part of a human studies research protocol which was approved by the Stanford University Institutional Review Board. The experiment was performed using fresh muscle specimens, according to availability of the clinical procedures. Sample processing for cell analysis began within one hour of specimen isolation.

### Drug Treatments

The inducible CreER recombinase system was activated by injecting Tamoxifen (Sigma-Aldrich) at a dose of 20 mg/mL in corn oil. Mice received 2 mg Tamoxifen intraperitoneally daily for seven consecutive days. For in vivo ketone body treatment, mice were injected with 0.6 g/kg BHB and 0.6 g/kg acetoacetate daily for 7 days. Doses were distributed over 3 injections per day. For in vivo BHB treatment, mice were injected with 2.6 g/kg BHB in PBS daily for 7 days. Doses were distributed over 4 injections per day. Unless otherwise stated, BHB injections or ex vivo treatments were done using a racemic mixture of S-BHB and R-BHB. For in vivo HDAC inhibition, mice were injected with 10 mg/kg givinostat, 30 mg/kg droxinostat, 20 mg/kg entinostat, 30 mg/kg tacedinaline, or 30 mg/kg MC1568 once daily for 7 days. For in vivo p53 stabilization, mice were injected once daily with 20 mg/kg nutlin-3. Final blood BHB concentrations of mice that were fasted, fed a ketogenic diet, or injected with ketone bodies were recorded just before sacrifice between 10:00 a.m. and 11:00 a.m. using the FDA approved Keto Mojo blood glucose and ketone monitoring kit. This kit uses an enzymatic BHB detection method that measures blood levels of R-BHB.

### Dietary Intervention

For ketogenic diet experiments, mice were given a diet (Bio-Serv S3666) that contained 7.24 kcal/g composed of 8.6% protein, 75.1% fat, and 3.2% carbohydrates. Control and refed mice were given 24-hour access to regular chow diet that contained 3.1 kcal/g composed of 18% protein, 6.2% fat, and 44.2% carbohydrates (Envigo Teklad 2018).

### Muscle Injury

For muscle injuries, mice were anesthetized with isoflurane before TA muscles were disinfected with ethanol and iodopovidone, injected with 50 µl BaCl_2_ (Sigma), and poked 50 times with a 31-gauge insulin syringe. Mice received post-surgery buprenorphine analgesia.

## METHOD DETAILS

### MuSC Isolation

MuSCs were isolated from mouse hindlimbs as previously described (Liu Ling, Cheung TH, 2016). Briefly, hindlimb muscle was dissected and homogenized manually with dissection tools before incubation with type II collagenase and Dispase digestive enzymes. Following enzymatic digestion, samples were mechanically dissociated using a 20-gauge needle and syringe then filtered through a 40 µm cell strainer. The bulk tissue prep was incubated with fluorescently labeled antibodies at 4°C for 1 hour before a subsequent filtering step through a 35 µm strainer and MuSC isolation by FACS. MuSCs were purified from bulk tissue prep by positive selection for cells expressing VCAM and negative selection against cells expressing CD45, CD31, and Sca1. Human MuSCs were purified from a fresh operative sample as previously described (Charville et al., 2015). The operative sample was carefully dissected from adipose and fibrotic tissue and a disassociated muscle suspension prepared as described for mouse tissue. The resulting cell suspension was then washed, filtered and stained with anti-CD31-Alexa Fluor 488 (clone WM59; BioLegend; #303110, 1:75), anti-CD45-Alexa Fluor 488 (clone HI30; Invitrogen; #MHCD4520, 1:75), anti-CD34-FITC (clone 581; BioLegend; #343503, 1:75), anti-CD29-APC (clone TS2/16; BioLegend; #303008, 1:75) and anti-NCAM-Biotin (clone HCD56; BioLegend; #318319, 1:75). Unbound primary antibodies were then washed, and the cells were incubated for 15 minutes at 4°C in streptavidin-PE/Cy7 (BioLegend) to detect NCAM-biotin. Cell sorting was performed to obtain the MuSC population as previously described (Charville et al., 2015).

### MuSC Culture

Freshly isolated MuSCs were plated in wells that were pre-coated with poly-D-lysine (0.1 mg/mL, EMD Millipore) and ECM (25 µg/mL, Sigma). Unless otherwise stated, cells were cultured in growth medium (GM: Ham’s F-10, 20% FBS, 2.5 ng/ml FGF, 100 U/mL penicillin, and 100 μg/mL streptomycin). To isolate cultured MuSCs for analyses, cells were treated with trypsin then pelleted by centrifugation at 1000*g* for 5 minutes.

### Single Muscle Fiber Isolation

Single myofibers were isolated as described (Rosenblatt et al., 1995). Briefly, extensor digitorum longus (EDL) muscles were dissected and digested by gentle agitation at 37°C in Ham’s F10 with 10% horse serum and 2 mg/ml collagenase II for 80 minutes. Digested muscle was triturated in Ham’s F10 with 10% horse serum using a wide bore glass pipet and a 10 cm culture dish. Isolated fibers were washed three times in medium before fixing.

### S Phase Entry

To monitor MuSC activation in vivo during homeostasis, mice received 50 mg/kg EdU via intraperitoneal injection daily for 2 weeks before MuSCs were isolated, fixed in 4% PFA, and stained for EdU. To monitor in vivo activation after injury, mice received 50 mg/kg EdU via intraperitoneal injection once at 60 hours post injury. Mice were then sacrificed at 72 hours post injury and MuSCs were isolated as described. Ex vivo S phase entry was monitored by supplementing the media with 10 µM EdU and staining using Click-iT™ EdU Cell Proliferation Kit (Invitrogen).

### Histology and Immunocytochemistry

For transplantation histology, TA muscles were fixed in 0.5% PFA and dehydrated in 20% sucrose prior to cryopreservation. For muscle regeneration histology muscles were frozen immediately after dissection. Cryopreserved TA muscles were sectioned at the belly of the muscle into 10 µm sections. Sections were then fixed in 2% PFA at room temperature for 10 minutes before staining. Sections were stained and imaged on a Zeiss Observer Z1 fluorescent microscope equipped with a Hamamatsu Orca-ER camera. Regenerating central nucleated muscle fiber size was quantified using the contour functions (findContours, contourArea, and arcLength) of the open source Python package, OpenCV (https://pypi.org/project/opencv-python/). The number of RFP^+^ muscle fibers per TA cross section was quantified by hand.

### MuSC Flow Cytometry Analysis

Mean forward scatter (FSC-A) of freshly isolated MuSCs was recorded to measure cell size. To measure RNA content, MuSCs were blocked for 45 minutes at 37°C in wash media (WM: 90% Ham’s F-10 + 10% horse serum) with 10 µM Hoechst 33342 (Thermo Fisher). Cells were then stained with 0.1 mg/mL Pyronin Y (Santa Cruz sc-203755) in wash media for 15 minutes at 37°C and analyzed on an Aria III. RNA content of each sample was recorded as mean fluorescence intensity in the PE-Cy5 channel. To measure mitochondrial content, MuSCs were incubated in 300 nM Mitotracker Green FM (Invitrogen M-7514) in wash media at 37°C for 45 minutes then analyzed on an Aria III. Mitochondrial content of each sample was recorded as mean fluorescence intensity in the FITC channel. Cell survival was measured by FACS analysis after staining cells with a membrane-impermeable DNA binding dye (DAPI, PI, or 7-AAD) and/or annexin V (BD Pharmingen).

### Proteomics

Shotgun proteomic samples were prepared by precipitating MuSC proteomes using 100% TCA (Sigma T6399), which was added to yield a final concentration of 20%. Samples were incubated at -80°C overnight to precipitate proteins and then centrifuged at 4°C for 10 minutes at 10,000*g*. The pellet was washed 3 times with ice cold 0.1 M HCl in 90% acetone, air-dried, and then resuspended in 30 µL 8M urea in PBS. ProteaseMAX™ (30 µL of 0.2% in 100 mM ammonium bicarbonate) was then added to samples, after which the samples were vortexed and diluted with 40 µL ammonium bicarbonate. TCEP was added to a final concentration of 10 mM and then samples were incubated for 30 minutes at 60°C, followed by addition of iodoacetamide to a final concentration of 12.5 mM, and samples were incubated at room temp for 30 minutes. Samples were diluted with 100 µL of PBS and 1.2 µL of 1% ProteaseMAX™ was added and vortexed well, after which 2 µg of sequencing trypsin was added and samples incubated overnight at 37°C. Trypsinized samples were acidified with 5% formic acid and centrifuged at 13200 rpm for 30 minutes. Purified peptides were pressure-loaded onto a 250 mm silica capillary tubing filled with 4 cm of Aqua C18 reverse-phase resin (Phenomenex # 04A-4299). The samples were then attached using a MicroTee PEEK 360 µm fitting (Thermo Fisher Scientific #p-888) to a 10 cm laser pulled column of 100 mm fused silica capillary packed with 10 cm Aqua C18 reverse-phase resin. Samples were subsequently analyzed using an Orbitrap Q Exactive Plus mass spectrometer (Thermo Fisher Scientific). Data was collected in data-dependent acquisition mode with dynamic exclusion enabled (60 seconds). One full MS (MS1) scan (400-1800 m/z) was followed by 15 MS2 scans (ITMS) of the nth most abundant ions. Heated capillary temperature was set to 200°C and the nanospray voltage was set to 2.75 kV. The samples were run using a two-hour gradient from 5% to 80% acetonitrile with 0.1% formic acid at 100 nl/minute. Data were extracted in the form of MS1 and MS2 files using Raw Extractor 1.9.9.2 (Scripps Research Institute) and searched against the Uniprot mouse database using ProLuCID search methodology in IP2 v.3 (Integrated Proteomics Applications, Inc) (Xu et al., 2015).

### RNA Sequencing

Freshly isolated MuSCs from individual mice were frozen in liquid nitrogen, and RNA was extracted with using Nucleospin RNA XS kit (Machery-Nagel). RNA was reverse transcribed using oligo(dT) primers with the SMARTer Ultra Low Input system (Takara). The cDNA was then sheared with a Covaris S2 ultrasonicator. End repair, multiplexed adapter ligation and 13–15 cycles of library amplification were performed using the Ovation Ultralow Multiplex system (NuGEN). Libraries underwent single-end 150-bp sequencing at the Stanford Genomics Facility with an Illumina NextSeq 500 to a depth of 20–40 million reads.

RNA-seq data processing and gene expression analysis were carried out as described (Brett et al., 2020). To process raw RNA-seq data, trim_galore was used for adapter and quality trimming (https://www.bioinformatics.babraham.ac.uk/projects/trim_galore)(quality cutoff 20, adapter stringency 1, final length filter 50). STAR (mismatch cutoff 4% of read length, no non-canonical junction alignments) was used to map reads to mm10 (Ensembl release 89, no patches) using transcript annotations from GENCODE. The featureCounts module of the Subread package was used to summarize unique mappings over genes.

For analysis of gene expression, genes lacking FPKM of at least 1.5 in at least three samples and genes lacking an Entrez ID were filtered out. Raw count data were then normalized for library preparation and sample collection batch effects using RUVs (variation factors *k* = 6). Differential gene-expression analysis was performed for genes with a normalized FPKM value of at least 6 in at least three samples using edgeR, with Cox–Reid estimations of tagwise dispersions and negative binomial GLM likelihood ratio tests. The Benjamini–Hochberg FDR control for multiple-hypothesis testing was used to produce *q* values. GSEA for coherent biological processes was performed using the MSigDB Hallmark gene sets (enrichment statistic *P* =1, ranking metric Signal2Noise, expressed gene set size range 15–500, which excluded only Hallmark PANCREAS BETA CELLS). FDR *q* values were calculated by gene permutation.

### Western Blotting

One hundred fifty thousand MuSCs were isolated from the lower hindlimb muscles of each mouse. Cells were pooled into pellets of 300,000 cells, with cells from two specimens in each pellet. Protein from each pellet was extracted in 100 µl of 2% SDS, 100 mM Tris, pH 7.4. Protein extracts were electrophoresed on a 4%–15% gradient polyacrylamide gel and then transferred to a PVDF membrane. The intensities of protein bands and background intensities were quantified by densitometry using ImageJ.

### RT-qPCR

One hundred thousand cells were collected and washed twice with PBS. RNA was extracted using an RNeasy Mini Kit (QIAGEN) according to the manufacturer’s instructions and reverse-transcribed using the High Capacity cDNA Reverse Transcription Kit (Invitrogen). An ABI 7900HT Fast Real-Time PCR system and custom synthesized primers (Invitrogen) were used to amplify cDNA of target genes by quantitative PCR. Target transcript levels relative to GAPDH transcript levels were quantified using the comparative CT method (Pfaffl, 2001). Measurements were taken in triplicates for each of three different experiments.

### Transplantation

Recipient NSG mice were prepared for transplantation with 1800 cGy of irradiation to the lower hindlimbs while under ketamine anesthesia followed by BaCl_2_ injury to the TA muscles as described 3 days prior to transplantation. For competitive transplantation, 10,000 freshly isolated MuSCs from each donor group were pooled into an Eppendorf tube prior to centrifugation at 1000*g* for 5 minutes. For non-competitive transplantation, 10,000 cultured cells were transferred to an Eppendorf tube then pelleted by centrifugation at 1000*g* for 5 minutes. Pelleted cells were resuspended in 30 µl of PBS and injected into the TA muscles of recipient mice that were under isoflurane anesthesia.

### Bioenergetics

MuSC oxygen consumption rate (OCR) was determined using a Seahorse XFp analyzer (Agilent). Cells were FACS-isolated as described, plated in an ECM-coated 96-well SeaHorse plate at 200,000 cells per well, and analyzed in Seahorse base medium (Agilent 103335-100 supplemented with 25 mM glucose, 2 mM glutamine, and 1 mM sodium pyruvate) 1 hour after plating.

### Quantitative Mass Spectrometry of BHB levels in MuSCs and Whole Muscle

MuSCs from fasted and ad lib fed mice were isolated as described above with the additional inclusion of 10 µM AZD3965 and 30µM syrosingopine (inhibitors of MCT1 and MCT4 respectively, which are the two BHB transporters expressed in MuSCs.) Following FACS isolation, MuSC pellets were flash frozen and stored at −80°C.

The polar metabolite fraction from frozen cell pellets was extracted in 180 μL of 40:40:20 acetonitrile/methanol/water with inclusion of an internal standard 13C-BHB (10 nmol). Following 30 seconds of thorough vortexing and 1 minute of bath sonication, the polar metabolite fraction (supernatant) was isolated by centrifugation at 13,000*g* for 15 minutes. Twenty microliters of this supernatant were analyzed by SRM-based targeted LC–MS/MS. For whole muscle, muscle was dounce homogenized in 500 µl of 40:40:20 acetonitrile/methanol/water with inclusion of an internal standard ^13^C-BHB (10 nmol). Following dounce homogenization, the polar metabolite fraction (supernatant) was isolated by centrifugation at 13,000*g* for 15 minutes. Twenty microliters of this supernatant were analyzed by SRM-based targeted LC–MS/MS. For separation of polar metabolites, normal-phase chromatography was performed with a Luna-5 mm NH2 column (50 mm × 4.60 mm, Phenomenex). Mobile phases were as follows: Buffer A, acetonitrile; Buffer B, 95:5 water/acetonitrile with 0.1% formic acid or 0.2% ammonium hydroxide with 50 mM ammonium acetate for positive and negative ionization mode, respectively. The flow rate for each run started at 0.2 mL/minute for 5 minutes, followed by a gradient starting at 0% B and increasing linearly to 100% B over the course of 45 minutes with a flow rate of 0.7 mL/minute, followed by an isocratic gradient of 100% B for 17 minutes at 0.7 mL/minute before equilibrating for 8 minutes at 0% B with a flow rate of 0.7 mL/minute. MS analysis was performed with an electrospray ionization (ESI) source on an Agilent 6430 QQQ LC–MS/MS. The capillary voltage was set to 3.0 kV, and the fragmentor voltage was set to 100 V. The drying gas temperature was 350°C, the drying gas flow rate was 10 L/minute, and the nebulizer pressure was 35 psi. BHB abundance was determined by SRM of the transition from precursor to product ion. For BHB detection, the optimum transition was determined to be from precursor of m/z 103 to product of m/z 59 in negative ionization mode at a collision energy of 5. BHB abundance was quantified by integrating the area under the peak and was normalized to the area under the curve of an isotopic ^13^C BHB internal standard value.

### Statistical Analysis

Statistical analyses were performed using GraphPad Prism 5 (GraphPad Software). Unless otherwise stated, significance was calculated using two-tailed, unpaired Student’s t-tests. Statistical significance for competitive transplantation was calculated with GraphPad Prism 5 using a two-tailed paired Student’s t-test for relative fold change. Differences were considered to be statistically significant at the p < 0.05 level (*p < 0.05, **p < 0.01, ***p < 0.001, ns: not significant). Unless otherwise noted, all error bars represent SEM. Sample size for each experiment is indicated in the corresponding figure legend.

## ACKNOWLEDGEMENTS

Thank you to all of the Rando lab members for helpful insight and/or technical assistance, especially Heather Ishak and Yip Yu for being excellent managers and making available the resources we needed to carry out our studies. We are also grateful to Peter Crawford for providing *OXCT1*^fl/fl^ mice. This work was supported by a training grant from the NIH (1 T32 GM 119995-1 A1) to the Stanford Stem Cell Biology and Regenerative Medicine PhD Program, a training grant from the Buck Institute for Research on Aging (T32AG000266-21) to D.I.B., and grants from the Department of Veterans Affairs (BLR&D and RR&D Merit Reviews) and the NIH (P01 AG036695, R37 AG023806, and R01 AR073248) to T.A.R. Images used in **Figure 4E** were created with BioRender.com

## AUTHOR CONTRIBUTIONS

D.I.B. and P.B. designed the studies and carried out experiments with assistance from J.H.T., C.W.N., J.S.B., A.D.M., S.K., L.L., H.D., J.K., L.A.M., S.M.L, and D.K.N. T.A.R. provided guidance throughout. D.I.B. and P.B. interpreted the results with guidance and input from T.A.R. D.I.B., P.B., and T.A.R. wrote the manuscript and assembled the data with assistance from C.N. and L.F.

## DATA AVAILABILITY

The data that support the findings of this study are available from the corresponding author upon request. RNA-Seq data have been deposited in the NCBI Gene Expression Omnibus with the accession code GSE184821.

---

**Figure S1.**
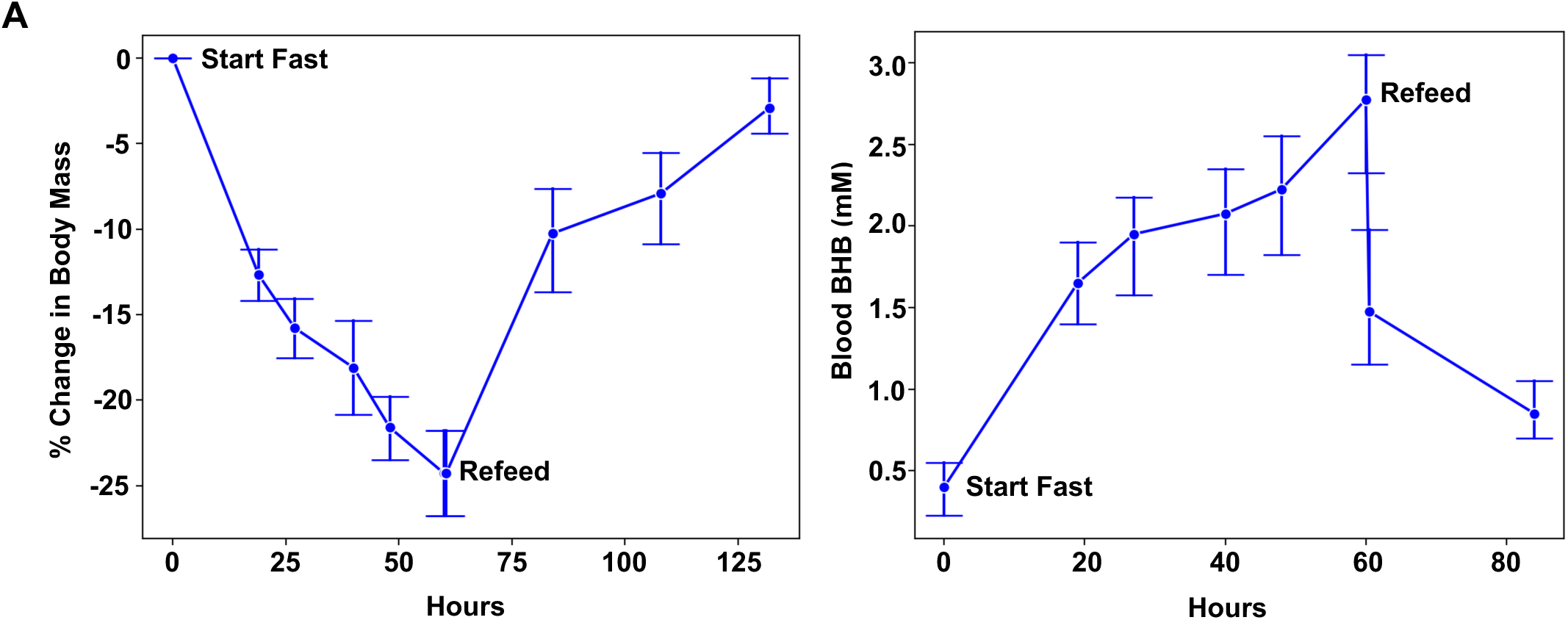
Fasting delays muscle regeneration, related to Figure 1. **A**) Percent change in body mass (left) and change in blood BHB concentration (right) of mice during fasting and after refeeding. Error bars represent SEM.

**Figure S2.**
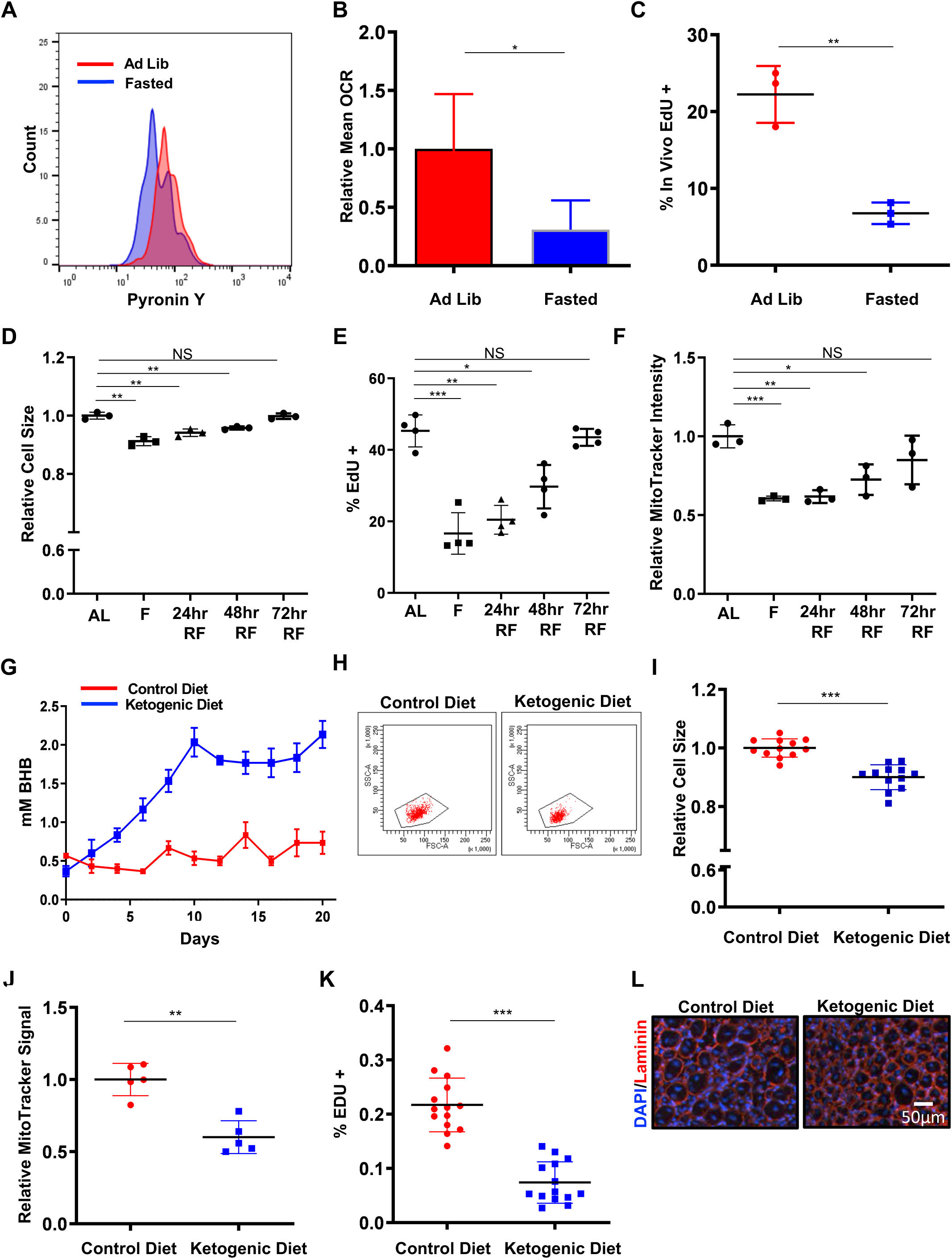
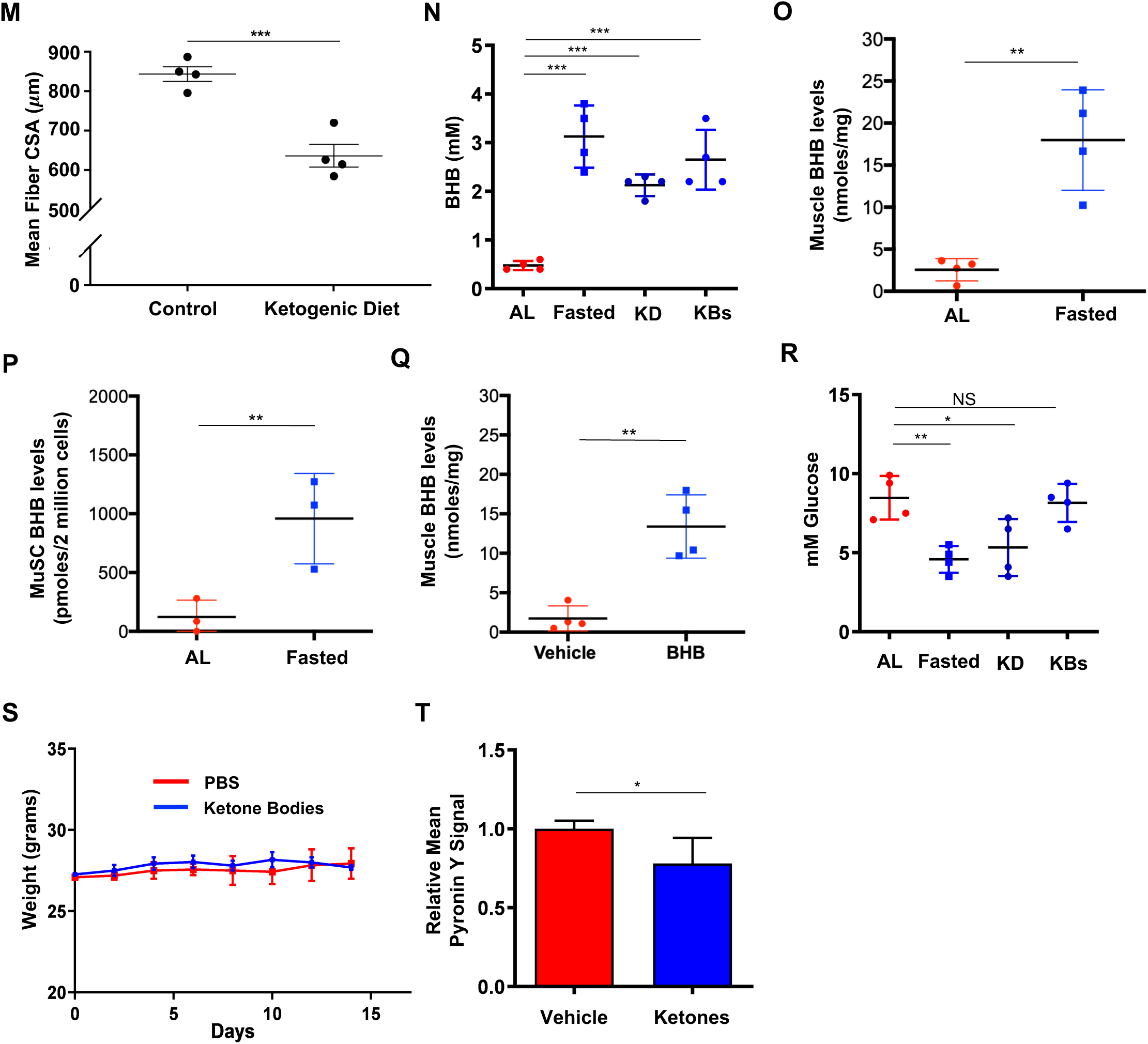
Fasting, a ketogenic diet, and exogenous ketosis can promote MuSC DQ, related to Figure 2. **A**) Representative FACS plot of Pyronin Y staining in freshly isolated MuSCs from fasted (60 hours) or ad lib fed mice. **B**) Quantification of relative basal oxygen consumption rates in freshly isolated MuSCs from fasted (60 hours) or ad lib fed mice (n=3). **C**) Quantification of in vivo EdU incorporation in MuSCs isolated from fasted (60 hours) and ad lib fed mice whose TA muscles had been injured with BaCl_2_ (n=3). **D**) Quantification of the cell size (based on forward scatter in FACS plots) of freshly isolated MuSCs from mice that were ad lib fed (AL), fasted (F) for 60 hours, or fasted for 60 hours and then refed (RF) for the indicated amount of time (n=3). **E**) Quantification of EdU incorporation in MuSCs isolated from mice as in panel (D). Cells from all groups were cultured for 48 hours in the continuous presence of EdU (n=4). **F**) Quantification of MitoTracker green signal intensity in freshly isolated MuSCs from mice as in panel (D) (n=3). **G**) Quantification of blood BHB levels in mice that were fed a ketogenic diet (3 weeks) or a control diet (n=4). **H**) Representative FACS plots of freshly isolated MuSCs from mice that were fed a ketogenic diet (3 weeks) or control diet. **I**) Quantification of the cell size (based on forward scatter in FACS plots) of freshly isolated MuSCs from mice that were fed a ketogenic diet (3 weeks) or control diet (n=12). **J**) Quantification of MitoTracker green signal intensity in freshly isolated MuSCs from mice that were fed a ketogenic diet (3 weeks) or a control diet (n=5). **K**) Quantification of EdU incorporation from MuSCs isolated from mice that were fed a ketogenic diet (3 weeks) or control diet. Cells from both groups were cultured for 48 hours in the continuous presence of EdU (n=14). **L**) Representative histology of regenerating muscle from mice fed a control or ketogenic diet (3 weeks) prior to BaCl_2_ injury to the TA muscle. TA muscles were harvested 7 days after injury, sectioned, and stained for laminin (red). Nuclei were stained with DAPI. **M**) Quantification of regenerating muscle fiber cross-sectional area from mice fed a control diet or a ketogenic diet (3 weeks) before injury as described in panel (L). **N**) Quantification of comparative blood BHB levels in mice after fasting (60 hours), ketogenic diet (3 weeks), or ketone body injections (200 mg/kg BHB and 200 mg/kg acetoacetate 3 times a day for 1 week) (n=4). **O-P**) Mass spec quantification of BHB levels in (O) TA muscles and (P) freshly isolated MuSCs from mice that were fed ad libitum or fasted for 60 hours. To prevent efflux of intracellular BHB during MuSC isolation, all samples were incubated with 10 µM AZD3965 and 30 µM syrosingopine (inhibitors of the BHB transporters MCT1 and MCT4 that are expressed in MuSCs) **Q**) Mass spec quantification of BHB levels in TA muscles 10 minutes after a bolus of 200 mg/kg BHB. **R**) Quantification of comparative blood glucose levels in mice after fasting (60 hours), ketogenic diet (3 weeks), or ketone body injections (1 week) (n=4). **S**) Quantification of body weights of mice after injections with ketone bodies or vehicle for two weeks. **T**) Quantification of relative mean Pyronin Y signal intensity in freshly isolated MuSCs from ketone body- or vehicle-treated mice (n=3). *p < 0.05; **p < 0.01; ***p < 0.001; ns, not significant.

**Figure S3.**
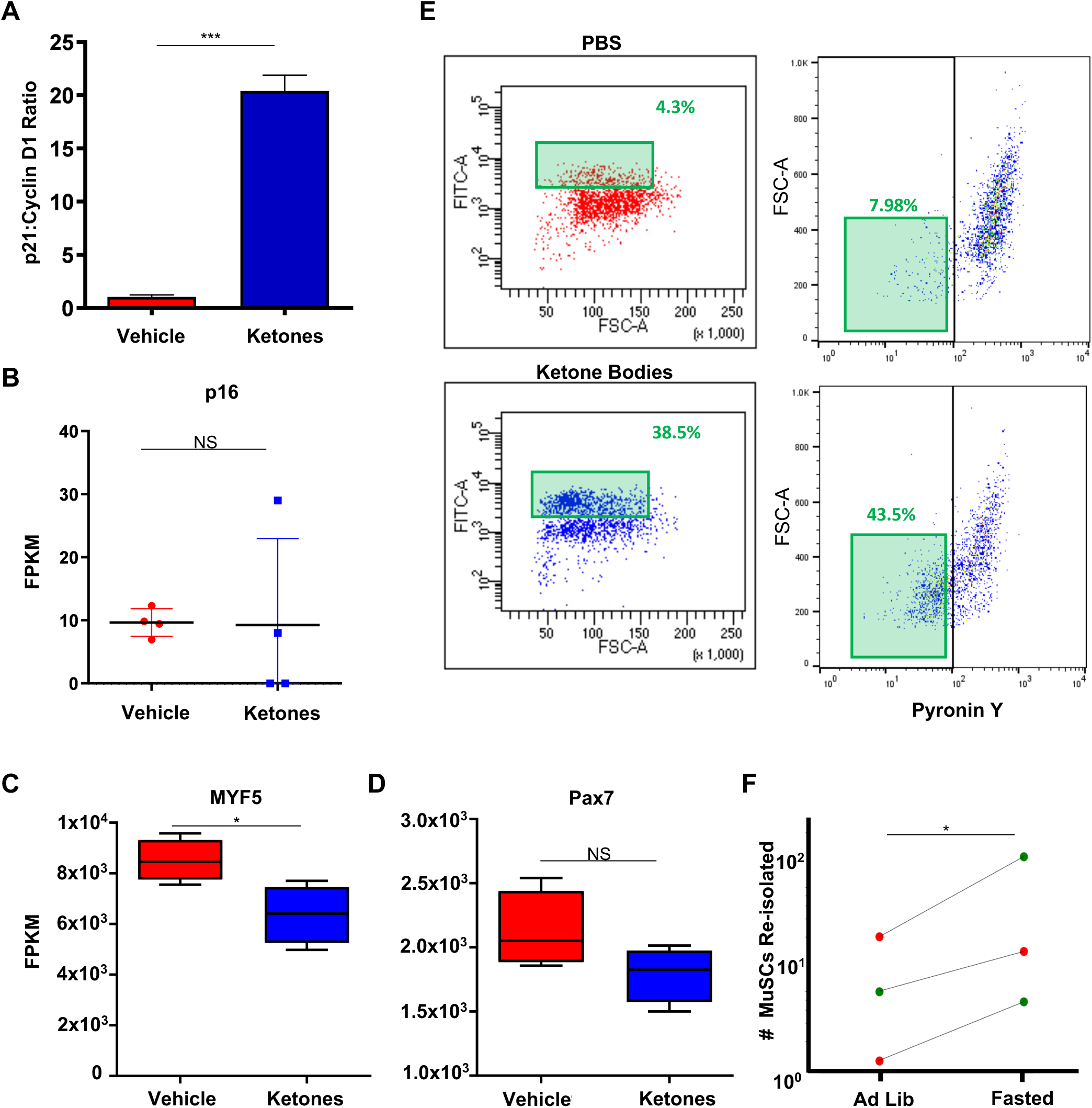
Ketosis promotes a transcriptional signature of DQ, related to Figure 3. **A**) Quantification of the ratio of p21 to cyclin D1 gene expression (from RNA-seq) in freshly isolated MuSCs from ketone body-treated or vehicle-treated mice (n=4). **B**) Quantification of p16 gene expression (from RNA-seq) in MuSCs from ketone body- or vehicle-treated mice (n=4). **C**) Quantification of MYF5 gene expression (from RNA-seq) in MuSCs from ketone body- or vehicle-treated mice (n=4). **D**) Quantification of Pax7 gene expression (from RNA-seq) in MuSCs from ketone body- or vehicle-treated mice (n=4). **E**) Representative FACS plots showing Pax7 promoter activity (FITC) and Pyronin levels in MuSCs from Tg(Pax7-ZsGreen) mice injected with ketone bodies or vehicle. MuSCs were plated in culture for 48 hours and analyzed by FACS for Pax7 promoter activity (left, top and bottom) and Pyronin Y levels (right, top and bottom). Highlighted with green boxes are the corresponding Pyronin Y levels of the Pax7^hi^ cells. **F**) Quantification of the number of MuSCs re-isolated from TA muscles of recipient mice 28 days after competitive transplantation. MuSCs isolated from fasted (60 hours) or ad lib fed mice with different color lineage tracers were pooled and transplanted into the same recipient muscle. Green data points represent the number of re-isolated donor cells with an eYFP Pax7 lineage tracer and red data points represent the number of re-isolated donor cells with a tdTomato Pax7 lineage tracer. Error bars represent SEM. *p < 0.05; **p < 0.01; ***p < 0.001; ns, not significant.

**Figure S4.**
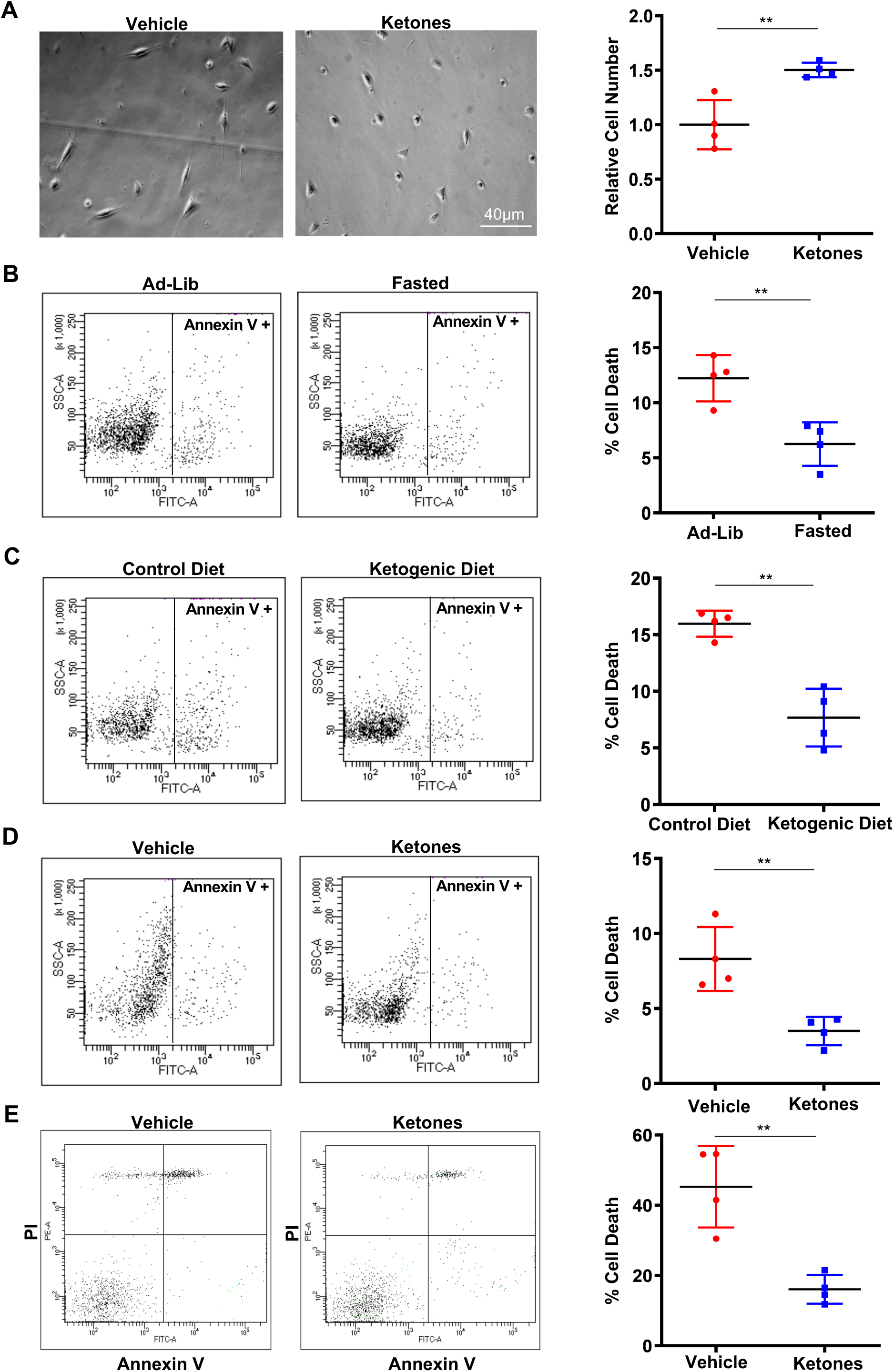
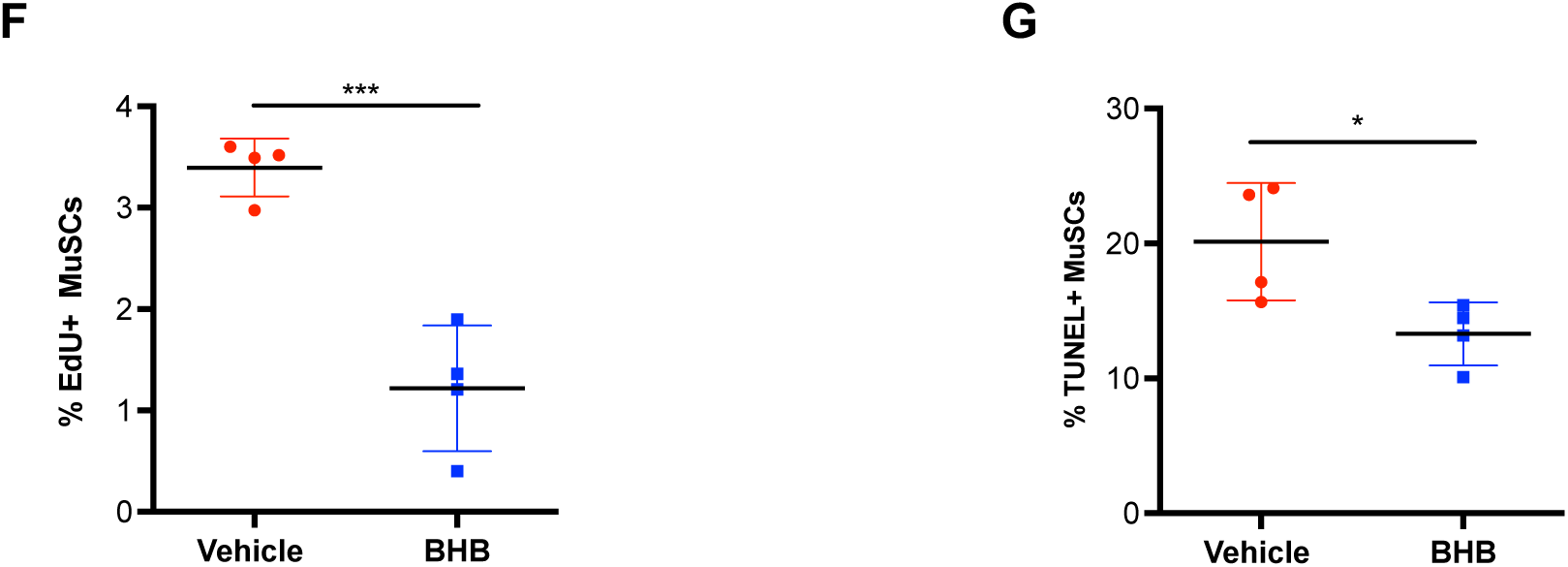
KIDQ promotes MuSC resilience, related to Figure 4. **A**) Representative brightfield images (left) and quantification (right) of the relative cell number of MuSCs (after 48 hours in culture) derived from ketone body- or vehicle-treated mice. Survival of MuSCs isolated from control mice or mice that were **B**) fasted for 60 hours, **C**) fed a ketogenic diet for 3 weeks, or **D**-**E**) injected with ketone bodies. MuSCs were probed for survival after 48 hours in culture (B-D) or after 8 hours in low nutrient media (E). Representative FACS plots (left) and quantification (right) of cell death by measuring the percentage of cells that were annexin V positive (B-D) or annexin V positive and PI positive (E). **F-G**) Quantification of (F) percent EdU incorporation and (G) percent cell death as measured by TUNEL staining in human MuSCs that were treated with vehicle or 5 mM BHB in growth media for 48 hours. Quantification of (n=4 for A-G). Error bars represent SEM. *p < 0.05; **p < 0.01; ***p < 0.001; ns, not significant.

**Figure S5.**
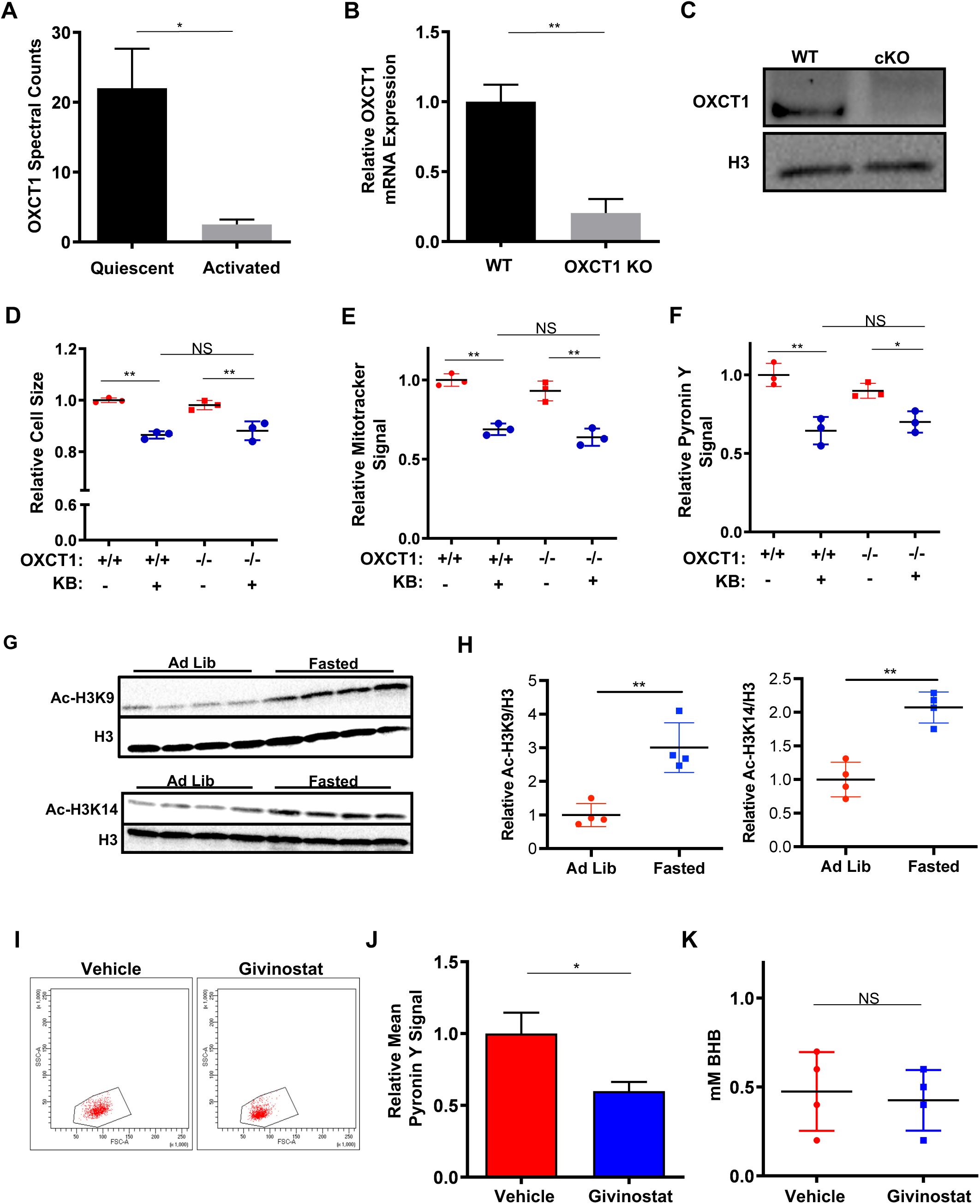
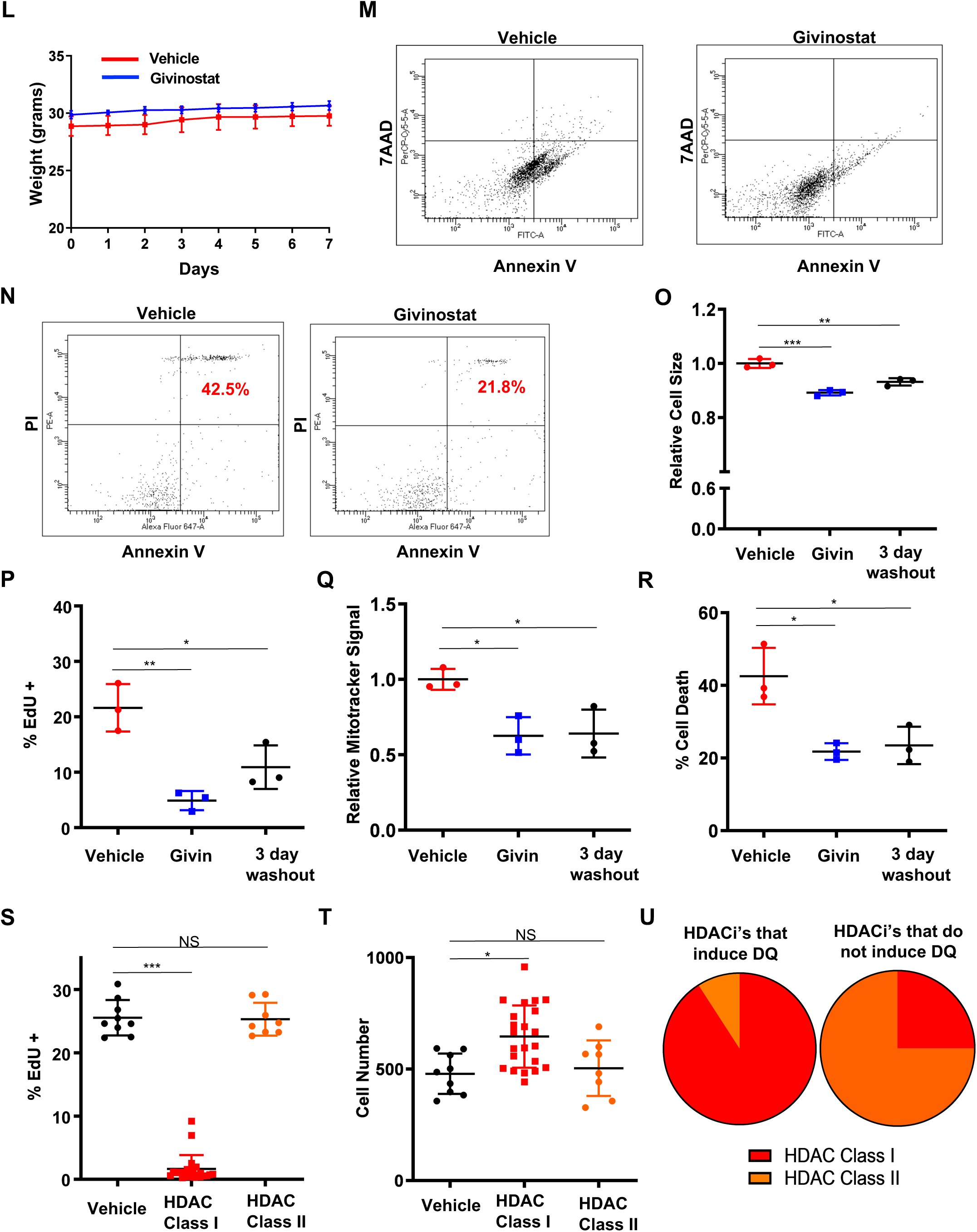
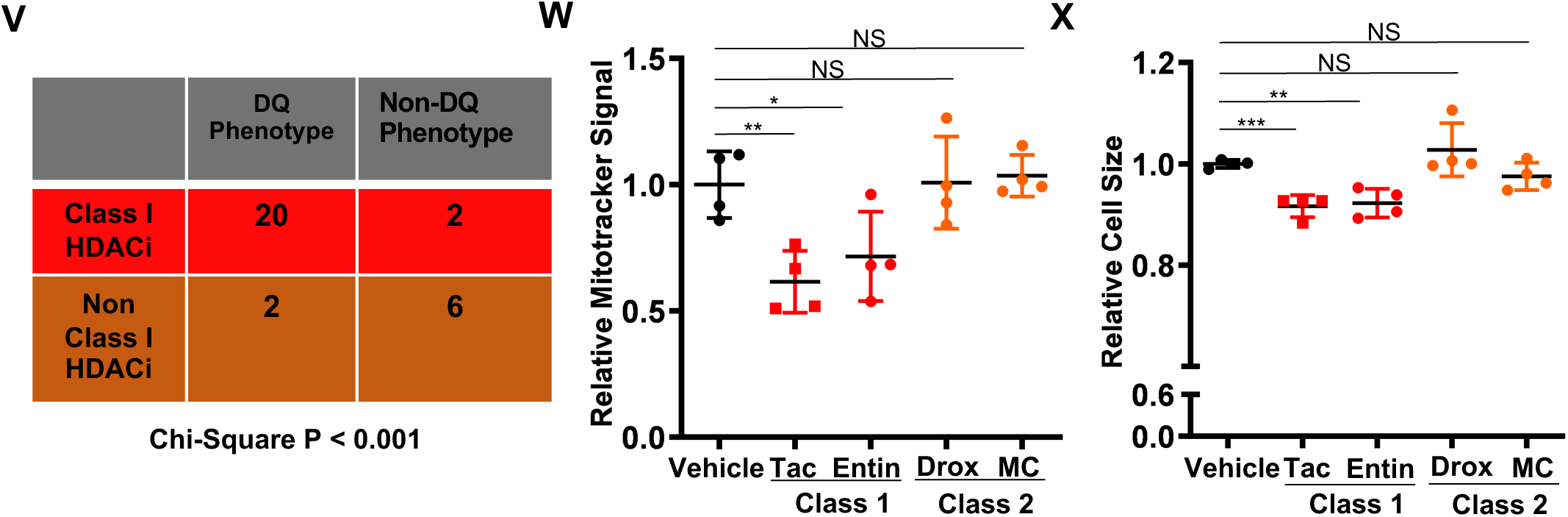
Ketone bodies elicit DQ via a nonmetabolic HDAC inhibitory mechanism, related to Figure 5. **A**) Quantification of OXCT1 protein spectral counts (from proteomics) between quiescent and 24 hour-activated MuSCs (n=2; 20 pooled mice per replicate). **B**) Quantification of relative OXCT1 mRNA expression in MuSCs derived from OXCT1 cKO or WT mice. Cells were isolated from uninjured muscle 14 days following tamoxifen treatment. **C**) Representative western blot analysis of OXCT1 protein levels in purified WT and p53 cKO MuSCs. Total H3 was used as a loading control. Cells were isolated from uninjured muscle 14 days following tamoxifen treatment. Quantification of **D**) relative cell size, **E**) relative MitoTracker intensity, and **F**) Pyronin Y signal intensity in freshly isolated MuSCs derived from OXCT1 cKO or WT mice following one week of in vivo treatment with either ketone bodies or vehicle (n=4). **G**) Western blots of H3K9 acetylation (top) and H3K14 acetylation (bottom) in freshly isolated MuSCs from mice that were fed ad libitum or fasted for 60 hours . Total H3 was used as a loading control **H**) Quantifications of Western blot analysis of H3K9 acetylation (left) and H3K14 acetylation (right) shown in (G). **I**) Representative FACS plots of cell size (forward scatter) and granularity (side scatter) and **J**) quantification of Pyronin Y signal intensity of freshly isolated MuSCs from mice injected with givinostat or vehicle. **K)** Quantification of blood BHB levels and **L**) body weights from mice injected with givinostat or vehicle. **M**-**N**) Representative FACS plots of MuSC survival from mice that were injected with givinostat or vehicle. MuSCs were probed for survival after (M) 48 hours in growth media by staining with annexin V and 7AAD and after (N) 8 hours in low nutrient medium (10% serum-free Ham’s F10 + 90% PBS) by staining for annexin V and propidium iodide. Numbers in the top right quadrant indicate the percentage of dead cells in each condition (n=4 for G-L). **O**-**R**) Mice were injected daily for one week with vehicle, givinostat, or givinostat followed by three days of vehicle. Depth of MuSC quiescence was determine based on (O) mean cell size at isolation (forward scatter), (P) percentage of EdU incorporation after 48 hours in culture, and (Q) MitoTracker signal intensity at isolation. (R) Death of freshly isolated MuSCs was measured by staining for annexin V and propidium iodide after 8 hours in low nutrient medium (n = 3 for M-P). **S**) Quantification of EdU incorporation and **T**) total cell number from WT MuSCs grown in culture for 48 hours with a panel of Class I and Class II-selective HDAC inhibitors in the continuous presence of EdU (n=8-22). **U**) Pie chart showing the different proportion of Class I and Class II-selective HDAC inhibitors that induce or do not induce DQ. Induction of DQ is determined by both the ability of an inhibitor to decrease S phase entry and increase cell number after 48 hours in culture. **V**) Table showing the number of different Class I and Class II-selective HDAC inhibitors that induce or do not induce DQ. **W**) Quantification of relative MitoTracker signal intensity and **X**) relative cell size (based on forward scatter in FACS plots) in freshly isolated MuSCs from mice treated with either the Class I-selective HDAC inhibitors (tacedenaline (tac) or entinostat (entin)), Class II-selective HDAC inhibitors (droxinostat (drox) or MC1568 (MC)), or vehicle for one week (n=4). Error bars represent SEM. *p < 0.05; **p < 0.01; ***p < 0.001; ns, not significant. Significance in (V) is determined by chi square test of independence.

**Figure S6.**
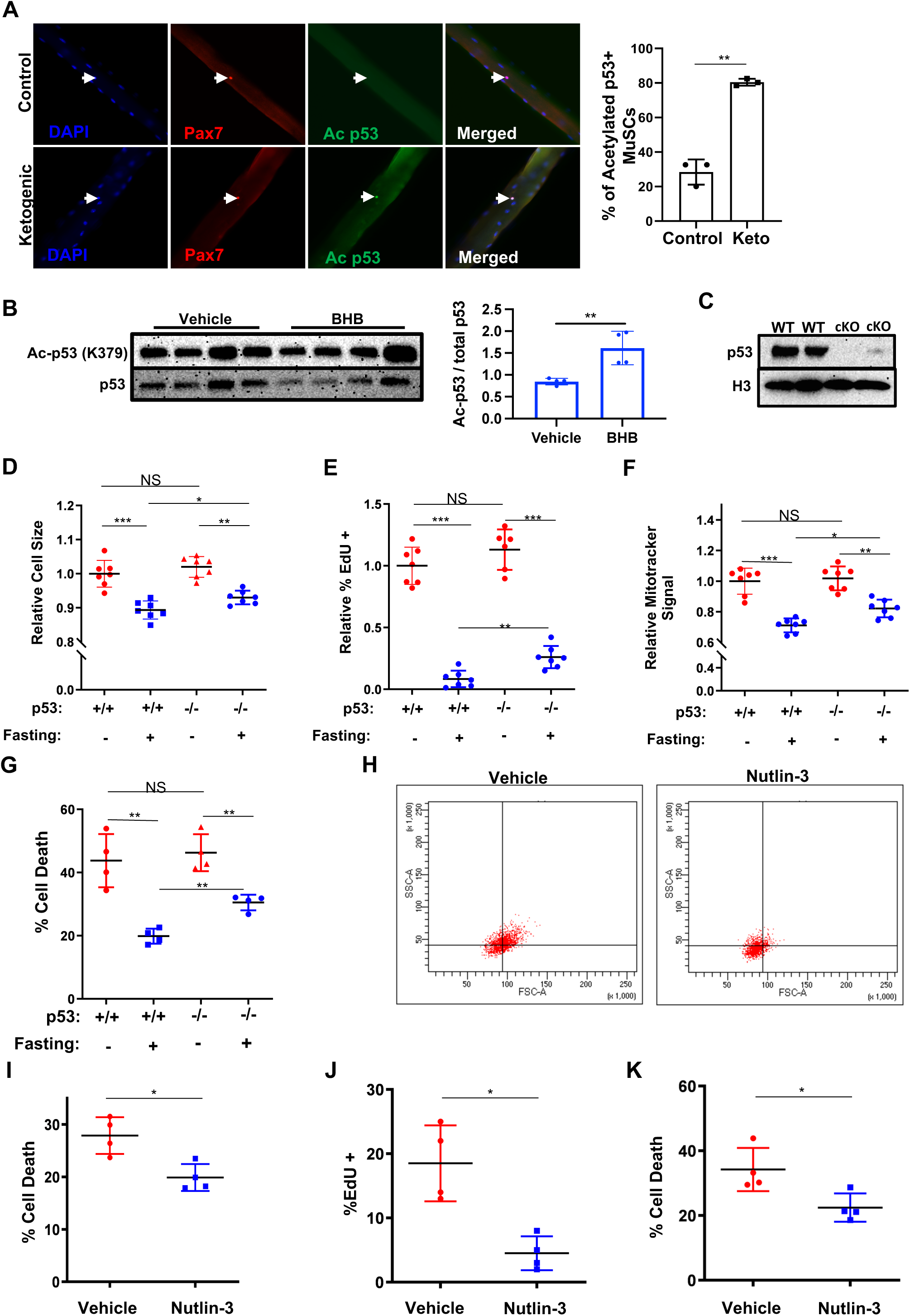
p53 is both necessary and sufficient to promote MuSC DQ, related to Figure 6. **A**) Immunofluorescence staining (left) for Pax7 and acetyl-p53 of single isolated myofibers taken from mice on a ketogenic diet (3 weeks) or control diet. Nuclei were stained with DAPI. Arrows indicate MuSCs. Quantification (right) of percent of Pax7+ MuSCs that stained positive for acetyl-p53 (n=3). **B**) Western blot analysis (left) and quantification (right) of p53 acetylation (K379) in MuSCs from mice injected with vehicle or BHB. Total p53 was used as a loading control for Ac-p53. **C)** Representative western blot analysis of p53 protein levels in purified WT and p53 cKO MuSCs. Total H3 was used as a loading control. Cells were isolated from uninjured muscle 14 days following tamoxifen treatment. **D**) Quantification of relative cell size in freshly isolated MuSCs derived from p53 cKO or WT mice that had been fasted (60 hours) or ad lib fed (n=7). **E**) Quantification of relative EdU incorporation in MuSCs derived from p53 cKO or WT mice that had been fasted (60 hours) or ad lib fed. Cells from all groups were maintained in culture for 48 hours in the continuous presence of EdU (n=6-7). **F**) Quantification of relative MitoTracker intensity in freshly isolated MuSCs derived from p53 cKO or WT mice that had been fasted (60 hours) or ad lib fed (n=7). **G**) Quantification of cell death in MuSCs derived from p53 cKO or WT mice that had been fasted (60 hours) or ad lib fed. Cell death was measured by cells that stained positive for both annexin V and propidium iodide following 8 hours of low-nutrient medium. **H**) Representative FACS plots of freshly isolated MuSCs from mice treated with the p53 activator nutlin-3a (20 mg/kg/day) or vehicle for one week. **I**) Quantification of cell death of freshly isolated MuSCs from mice treated with the p53 activator nutlin-3a (20 mg/kg/day) or vehicle for one week. Cell death was determined by cells that stained positive for both annexin V and propidium iodide (n=4). **J**) Quantification of EdU incorporation in WT MuSCs treated ex vivo with nutlin-3a (10 μM) or vehicle for 48 hours in the continuous presence of EdU (n=4). **K**) Quantification of cell death in WT MuSCs treated ex vivo with nutlin-3a (10 μM) or vehicle for 48 hours. After 48 hours in culture, cell death was determined by the percentage of cells that stained positive for both annexin V and propidium iodide (n=4). Error bars represent SEM. *p < 0.05; **p < 0.01; ***p < 0.001; ns, not significant.

